# Coupling of oxytocin and cholecystokinin pathways in the hypothalamus is required for gut-to-brain homeostatic feeding control

**DOI:** 10.1101/2022.07.20.500778

**Authors:** T Gruber, F Lechner, C Murat, RE Contreras, E Sanchez-Quant, V Miok, O Le Thuc, I González-García, RH Williams, PT Pfluger, TD Müller, SC Woods, CP Martinez-Jimenez, MH Tschöp, V Grinevich, C García-Cáceres

## Abstract

Oxytocin-expressing paraventricular hypothalamic neurons (PVN^OT^ neurons) integrate afferent signals from the gut including cholecystokinin (CCK) to adjust whole-body energy homeostasis. However, the molecular underpinnings by which PVN^OT^ neurons orchestrate gut-to-brain feeding control remain unclear. Here, we show that mice undergoing selective ablation of PVN^OT^ neurons fail to reduce food intake in response to CCK and develop hyperphagic obesity on chow diet. Notably, exposing wildtype mice to a high-fat/high-sugar (HFHS) diet recapitulates this insensitivity towards CCK, which is linked to diet-induced transcriptional and electrophysiological aberrations specifically in PVN^OT^ neurons. Restoring OT pathways in DIO mice via chemogenetics or polypharmacology sufficiently re-establishes CCK’s anorexigenic effects. Lastly, by single-cell profiling, we identify a specialized PVN^OT^ neuronal subpopulation with increased κ-opioid signaling under HFHS diet, which restrains their CCK-evoked activation. In sum, we here document a novel (patho)mechanism by which PVN^OT^ signaling uncouples a gut-brain satiation pathway under obesogenic conditions.

## Introduction

Despite historical evidence, as well as recent large-scale genetic studies (Locke et al. 2015) cogently linking obesity pathogenesis to central nervous system (CNS) defects, we still do not know the exact cellular and molecular mechanism(s) involved in the initiation and progression of obesity and related metabolic disorders. Multiple studies have highlighted various sets of hypothalamic neurons that release specific neuropeptides that critically govern energy intake versus expenditure. Among these, oxytocin (OT), a nine-amino acid neuropeptide traditionally recognized for its role in reproductive physiology and social behavior, is increasingly gaining attention as an anti-obesity drug due to its favorable metabolic effects in multiple pre-clinical and clinical studies (McCormack, Blevins, and Lawson 2020).

In the brain, endogenous OT is exclusively synthesized by neurons of the supraoptic (SON), accessory (AN) and paraventricular (PVN) nuclei of the hypothalamus (Swanson and Sawchenko 1983). Among these, the PVN has a particularly paramount role with regard to metabolic homeostasis as both electrolytic lesions as well as human genetic defects that impede PVN development result in severe obesity (Cox and Sims 1988; Michaud et al. 2001; Faivre et al. 2002; Tolson et al. 2010). In contrast to the evolutionarily more ancient, anatomically simpler SON and AN (Grinevich et al. 2014; Knobloch and Grinevich 2014), the PVN exhibits a more complex cytoarchitecture (Swanson and Sawchenko 1980; Biag et al. 2012) and harbors multiple cell types including two types of OT neurons: ‘magnocellular’ and ‘parvocellular’ neurons (magnOT and parvOT, respectively), which differ in size, morphology, electrophysiological properties, axonal projection targets, and other properties (Althammer and Grinevich 2017).

Several investigations have found that gastric distension upon meal ingestion, and particularly the associated release of the gut peptide cholecystokinin (CCK), powerfully stimulate electrical activity of certain subsets of OT neurons (Verbalis et al. 1986; Renaud et al. 1987; Leng, Way, and Dyball 1991; Kutlu et al. 2010; Caquineau, Douglas, and Leng 2010). Chronic exposure to hypercaloric diets and the consequent obesity lead to defects in this gut-brain crosstalk and these in turn have been proposed to further aggravate metabolic derailment (Clemmensen et al. 2017; Brandsma et al. 2015). Indeed, the food intake-suppressing effect of CCK is severely attenuated upon high-fat diet feeding, and it in turn is associated with reduced neural activation in several hypothalamic nuclei, including the PVN (French et al. 1995; Covasa 2010; Troy et al. 2016).

We first found that the hypothalamic OT system undergoes maladaptive changes during chronic overnutrition in mice, including blunting the integration and propagation of the afferent CCK signal at the level of the OT system; i.e., a compromised OT response in the PVN is paramount for the loss of CCK’s appetite suppression under high-fat/high-sugar (HFHS) diet feeding. We then, by taking advantage of various gain-and-loss-of-function models, systematically interrogated the physiological relevance and therapeutic potential of the OT system in obesity and diabetes pathogenesis. Last, we employed single-nucleus RNA sequencing-2 (snRNA-seq2) and identified significant disruptions in the transcriptional profiles of specific OT subpopulations upon exposure to a HFHS diet. In sum, we shed important new light on the hitherto elusive molecular underpinnings of how OT neurons influence CCK-induced hypophagia in the orchestration of whole-body metabolic homeostasis.

## Results

### Selective adult-onset PVN^OT^ neuron ablation induces rapid hyperphagic obesity

To assess if OT neurons located in the PVN (PVN^OT^ neurons) have a significant role in the control of energy homeostasis, we targeted these cells for selective ablation in mice using a diphteria toxin A (DTA)-based genetic approach. Therefore, we stereotaxically injected adult male mice that carry the DTA gene downstream of a <u>L</u>oxP-<u>S</u>TOP-<u>L</u>oxP (LSL) with adeno-associated viruses (AAV) that drive expression of iCre or the fluorescent reporter Venus under an OT promoter (Knobloch et al. 2012; Grinevich et al. 2016) (Figure 1A). As expected, iCre-mediated excision of the LSL cassette resulted in prominent induction of cleaved caspase 3 (C-CASP3; an apoptosis marker) in PVN^OT^ neurons at three days post AAV injection (Figure 1B). Importantly, while PVN^OT^ neurons were greatly reduced in number at the end of the experiment (8 weeks post AAV injection), neighboring neurons expressing arginine-vasopressin (AVP) (Figure S1A, S1B) as well as OT and AVP neurons of the SON and accessory nuclei remained unaffected (data not shown). At two weeks post stereotaxic surgery, DTA^OT+/PVN^ mice had already substantially and significantly increased body weight compared to their control littermates and eventually developed pronounced obesity on standard chow (SC) diet with age (Figure 1C, D).

**Figure 1:**
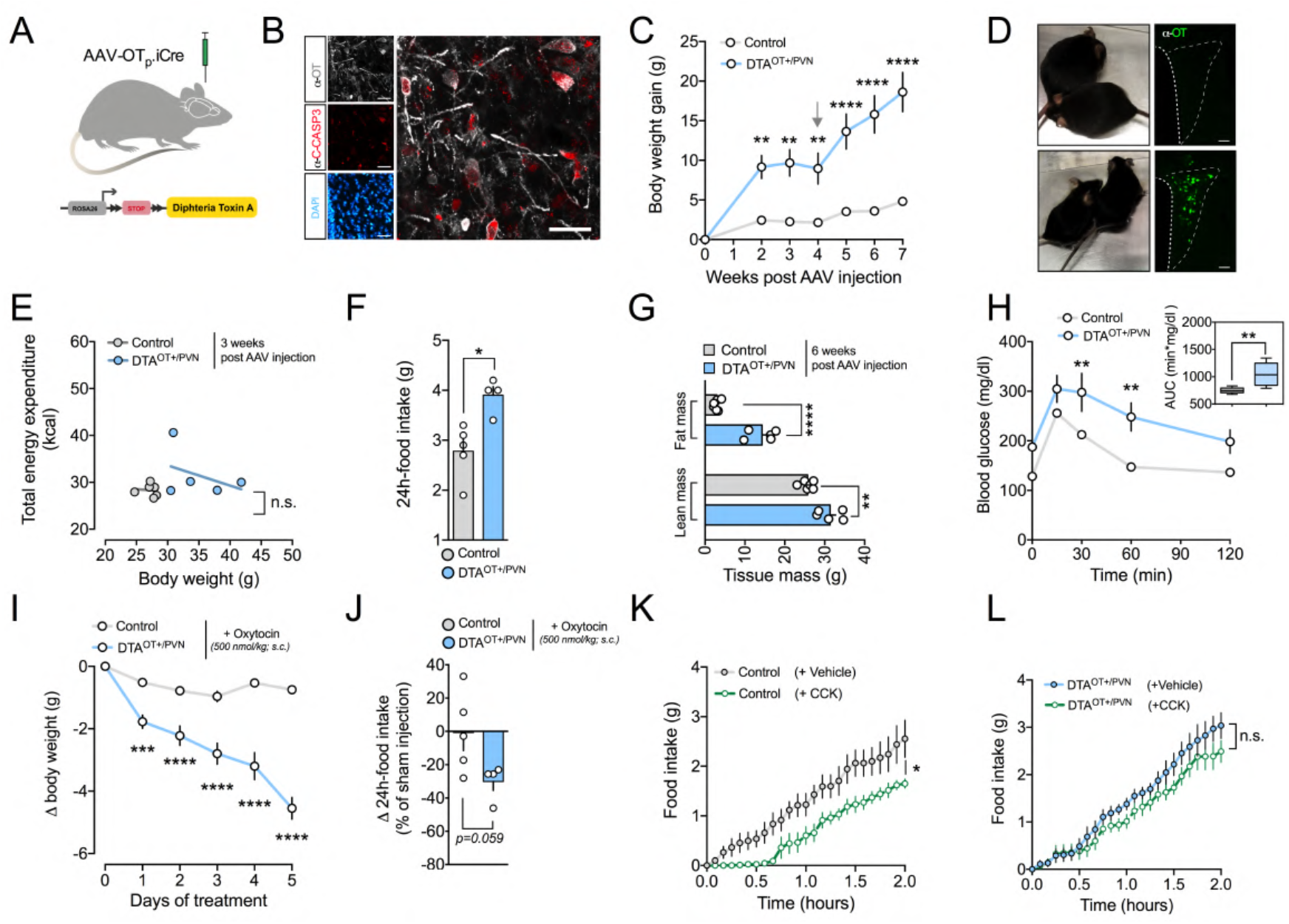
Virus-mediated ablation of PVN^OT^ neurons induces hyperphagic obesity that is rectifiable by exogenous oxytocin treatment and associated with CCK insensitivity. **(A)** Schematic illustration of the experimental paradigm in which an OT-specific AAV-OTp-iCre is used to induce the expression of diphtheria toxin A (DTA) selectively in PVN^OT^ neurons. **(B)** Representative confocal micrograph depicting immunoreactivity to cleaved caspase 3 (C-CASP3; red) in PVN^OT^ neurons (grey) five days post injection. Scale bar, 50 μm. **(C)** Body weight gain of DTA^OT+/PVN^ and control mice fed SC diet. Arrow at three weeks post AAV injection indicates single housing in metabolic cages. Data are presented as mean ± SEM. ** P < 0.01, **** P < 0.0001. n = 5-7 mice (two-way ANOVA). **(D)** Representative images of two pairs of DTA^OT+/PVN^ and control mice five weeks post AAV injection (upper and lower panel, respectively) next to epifluorescent micrographs depicting absence and presence of PVN^OT^ neurons, respectively. Scale bar, 100 μm. **(E)** Linear regression analysis of total energy expenditure (EE; assessed by indirect calorimetry) and body weight of DTA^OT+/PVN^ and control mice two weeks post AAV injection. Data are presented as individual mice. n.s., not significant. n = 5-7 mice. **(F)** Daily food intake of a separate cohort of DTA^OT+/PVN^ and control mice that were pair-housed in order to mitigate isolation stress (s. supplementary information). Data are presented as mean ± SEM. * P < 0.05. n = 4-5 pair of mice (unpaired Student’s *t*-test). **(G)** Body composition analysis of DTA^OT+/PVN^ and control mice six weeks post injection. Data are presented as mean ± SEM. ** P < 0.01, **** P < 0.0001. n = 5-7 mice (unpaired Student’s *t*-test). **(H)** Blood glucose changes of DTA^OT+/PVN^ and control mice upon a glucose tolerance test (2 g/kg BW *i*.*p*.; left panel) and area under the curve (right panel). Data are presented as mean ± SEM. ** P < 0.01, **** P < 0.0001. n = 5-7 mice (two-way ANOVA). **(I)** Body weight change in a separate cohort of DTA^OT+/PVN^ and control mice upon treatment with exogenous oxytocin (500 nM/kg BW *s*.*c*. twice daily). Data are presented as mean ± SEM. **** P < 0.0001. n = 8-10 mice (two-way ANOVA). **(J)** Change in food intake of pair-housed DTA^OT+/PVN^ and control mice upon treatment with exogenous oxytocin (500 nM/kg BW *s*.*c*. twice daily) relative to sham injections. Data are presented as mean ± SEM. **** P < 0.0001. n = 4-5 pairs of mice (unpaired Student’s *t*-test). **(K)** Cumulative food intake of control mice (nanoinjected with AAV-OTp-Venus) upon vehicle versus CCK (20 μg/kg BW *i*.*p*.). Data are presented as mean ± SEM. * P < 0.05, ** P < 0.01. n = 7 mice in a cross-over design (two-way ANOVA). **(L)** Cumulative food intake of DTA^PVN(OT+)^ mice (nanoinjected with AAV-OTp-iCre) upon vehicle versus CCK (20 μg/kg BW *i*.*p*.). Data are presented as mean ± SEM. n.s., not significant. n = 5 mice in a cross-over design (two-way ANOVA).

Indirect calorimetry conducted between the third and fourth week post AAV injection did not reveal differences between DTA^OT+/PVN^ mice and littermate controls with regard to uncorrected energy expenditure (Figure S1D), nor in the relationship between total energy expenditure and body weight (Figure 1E). Further, no change was observed in the respiratory exchange ratio (RER), in locomotor activity, or in cumulative food intake when single-housed in metabolic cages (Figure S1D-F). When pair-housed in their habitual home cages, however, a separate cohort of DTA^OT+/PVN^ mice displayed marked hyperphagia relative to littermate controls four weeks post AAV injection (Figure 1F), which led us to presume that differences in social isolation stress associated with the metabolic cages might have partly masked normal feeding behavior. Consistent with a persistent positive energy balance due to a higher food intake, DTA^OT+/PVN^ mice exhibited substantial differences in body composition at the end of the experiment (seven weeks post AAV injection), with significantly greater accrual of both fat mass and lean mass relative to littermate control mice (Figure 1G). Moreover, DTA^OT+/PVN^ mice had significantly impaired glucose tolerance (Figure 1H) and higher levels of glycated hemoglobin A_1C_ (HbA_1C_), implying a defect of long-term glycemic control relative to control mice (Figure S1G). Presupposing that these changes in DTA^OT+/PVN^ mice were due to diminished endogenous OT signaling, we asked whether pharmacologically resubstituting OT could normalize feeding and/or the metabolic derailments. While bi-daily administration of exogenous OT (500 nmol/kg BW; *s*.*c*.) did not significantly alter body weight in lean control mice, it led to a rapid body weight reduction in DTA^OT+/PVN^ mice (Figure 1I; treatment initiated eight weeks post AAV injection). Notably, changes in body weight were accompanied by reduced food intake (Figure 1K) and improved 3h-fasted blood glucose (Figure S1H), as well as in insulin sensitivity (HOMA-IR; Figure S1I). Intriguingly, we noted that the potency of exogenous OT in DTA^OT+/PVN^ mice greatly surpassed the weight-lowering effect that has been previously reported in diet-induced obese (DIO) C57BL/6J wildtype mice of similar adiposity using comparable OT dosing (Snider et al. 2019). To determine if this heightened sensitivity of DTA^OT+/PVN^ mice to exogenous OT is consequent of diminished endogenous production, we observed a compensatory up-regulation of *Otr* mRNA expression (encoding for the OT receptor) in the hypothalami of DTA^OT+/PVN^ mice relative to littermate controls (Figure S1J). In sum, the selective ablation of hypothalamic PVN^OT^ neurons promotes hyperphagic obesity, likely via a paucity of endogenous OT signaling that can be rectified by pharmacological substitution.

### Selective adult-onset PVN^OT^ neuron ablation renders mice insensitive to systemic CCK

PVN^OT^ neurons are strongly excited in response to meal-related gastrointestinal stimuli including gastric distension and particularly the associated release of the intestinal peptide CCK (Verbalis et al. 1986; Renaud et al. 1987; Leng, Way, and Dyball 1991; Kutlu et al. 2010; Caquineau, Douglas, and Leng 2010). Given that CCK administration robustly suppresses acute feeding, we assessed the necessity of PVN^OT^ neuron activation for CCK-elicited food intake suppression using mice devoid of PVN^OT^ neurons fed an SC diet (four weeks post AAV injection). After acclimating DTA^OT+/PVN^ mice and littermate controls to single housing, SC diet removal (3pm-6pm), and sham injections within metabolic cages for three days, we administered CCK (20 μg/kg BW; *i*.*p*.) on the fourth day to all mice 10 minutes before dark onset and food return. Consistent with the literature (Fan et al. 2004), CCK injections at this dose produced a significant suppression of food intake in control mice (Figure 1K). In contrast, however, feeding behavior of DTA^OT+/PVN^ mice was not significantly altered by CCK relative to sham treatments (Figure 1L). We conclude that PVN^OT^ neurons are necessary for CCK-mediated hypophagia under physiological conditions.

### Chronic exposure to a HFHS diet diminishes the activation of PVN^OT^ neurons to peripheral CCK

These findings suggest a prominent role of PVN^OT^ neurons in feeding control by CCK. We then asked whether this same gut-brain communication is altered by exposure to an obesogenic diet. Indeed, it has been reported that the food intake-suppressing effect of CCK is severely attenuated upon high-fat diet feeding, and this is associated with reduced neural activation in several hypothalamic nuclei including the PVN (French et al. 1995; Covasa 2010; Troy et al. 2016). Thus, we asked whether the hypothalamic OT system undergoes diet-induced desensitization to CCK upon HFHS diet feeding. Consistent with previous reports in rats (Olson et al. 1992; Motojima et al. 2016), C57BL/6J mice fed a SC diet and receiving CCK (20 μg/kg BW; *i*.*p*.) had robustly induced neuronal activation in a large proportion of PVN^OT^ neurons as indicated by increased co-localization with c-fos immunoreactivity (Figure 2A, B). To discern if peripheral CCK administration similarly affected both magnOT or parvOT subtypes, we next quantified c-fos^+^ PVN^OT^ neurons in a separate cohort of SC diet-fed reporter mice expressing tdTomato specifically in OT neurons (*OT:Ai14*), which received peripheral injections of Fluorogold (15 mg/kg BW; *i*.*p*.) 7 days prior to sacrifice. Fluorogold selectively marks neurons projecting beyond the blood-brain-barrier and thus can distinguish for magnOT cells (FG^+^; (Tang et al. 2020; Eliava et al. 2016). We found that 25% of CCK-activated PVN^OT^ neurons were FG^+^ magnOT neurons (Figure 2C, S2A-C), while CCK treatment induced c-fos only in 8% of FG^-^ parvOT neurons. Conversely, mice chronically fed an HFHS diet failed to significantly increase c-fos immunoreactivity in PVN^OT^ neurons following CCK injection suggesting a blunted activation in the course of consuming an obesogenic diet (Figure 2A, B).

**Figure 2:**
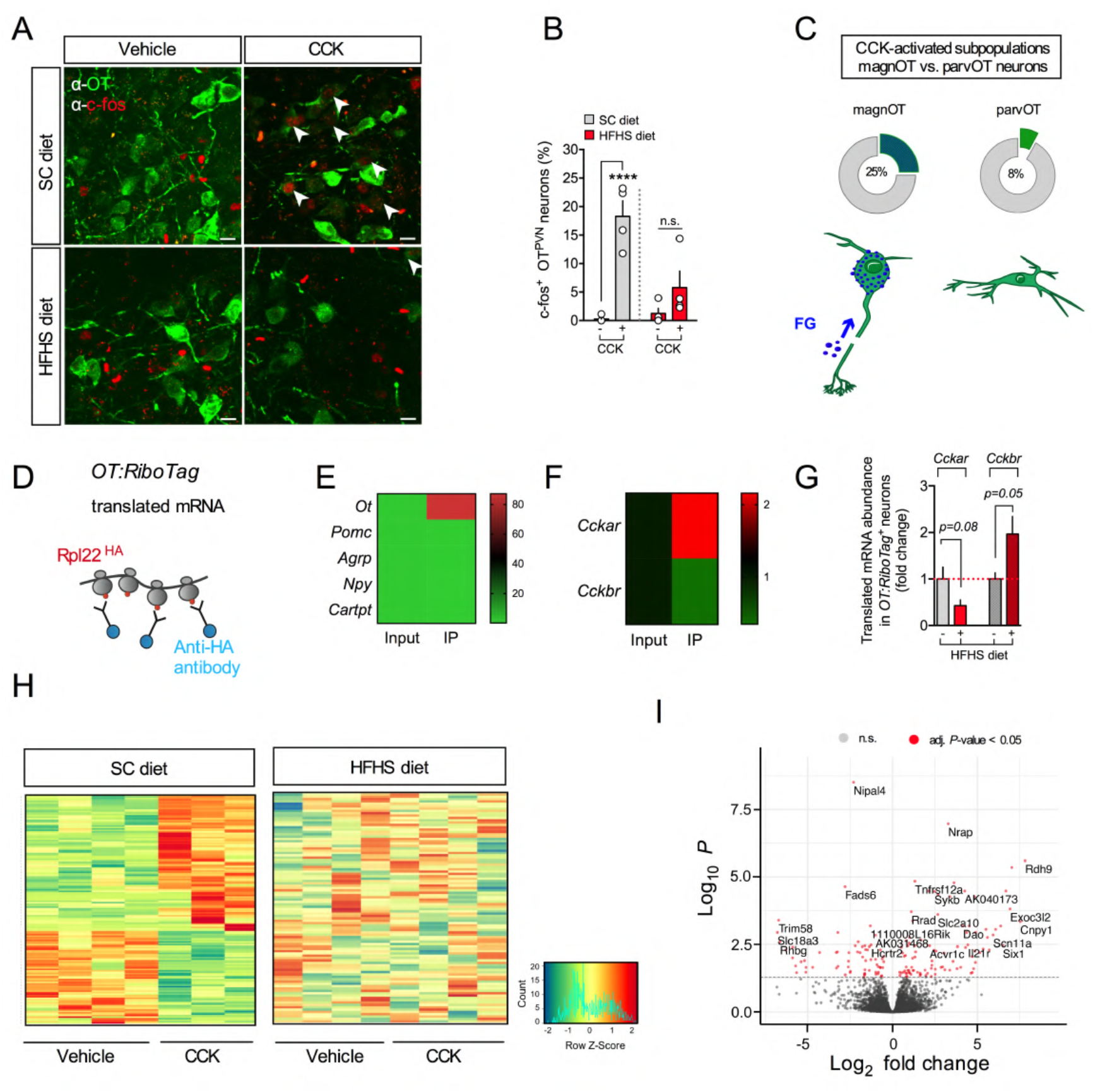
Chronic exposure to a HFHS diet impairs the electrical and transcriptional activation of PVN^OT^ neurons in response to peripheral CCK. **(A)** Representative confocal micrographs depicting neuronal activation by means of nuclear c-fos immunoreactivity (red) in PVN^OT^ neurons (green) in adult male C57BL/6J mice fed either SC or HFHS diet receiving CCK (20 μg/kg BW *i*.*p*.). Scale bar, 10 μm. **(B)** Corresponding quantification of c-fos^+^ PVN^OT^ neurons relative to total PVN^OT^ neurons counted. Data are presented as mean ± SEM. **** P < 0.0001, n.s., not significant. n = 4 mice, 4-8 hemisections per mouse (unpaired Student’s *t*-test). **(C)** Quantification of CCK-activated (c-fos^+^) subpopulations of PVN^OT^ in a separate cohort pre-treated with fluorogold (FG; 15 mg/kg BW *i*.*p*.) to distinguish parvOT neurons (FG^-^) from magnOT neurons (FG^+^); data is represented as mean in percent relative to total parvOT and magnOT cell count, respectively, and visualized as pie charts. n = 3 mice, 46 hemisections, 2486 cells. **(D)** Schematic illustration of the *OT:RiboTag* mouse model used to isolate actively translated mRNA specifically from OT^+^ neurons by immoprecipitation of HA-tagged ribosomal subunit Rpl22. **(E)** Heat map of translating mRNA enrichment of various hypothalamic neuropeptides in the immunoprecipitate (IP) relative to input. n = 4 mice. **(F)** Heat map representation of enrichment in *Cckar* mRNA and *Cckbr* mRNA in IP relative to input. N = 4 mice. **(G)** Relative abundance of *Cckar* mRNA and *Cckbr* mRNA in the IP of *OT:RiboTag* mice fed HFHS diet relative to SC diet. n = 4 mice. **(H)** Heat map representation of DEG under SC diet in hypothalamic OT^+^ neurons of adult male *OT:RiboTag* mice fed either SC (left) and HFHS (right) diet upon injection of CCK (20 μg/kg BW *i*.*p*.; tissue collection 2 h post-injection). Rows reflect normalized (Z score) gene expression abundance. n = 4 mice. **(I**) Volcano plot highlighting the gene expression changes, where the log transformed adjusted p-values are plotted against fold changes in hypothalamic OT^+^ neurons from OT:RiboTag mice fed HFHS diet relative to SC diet-fed control mice

### Peripheral CCK induces a characteristic transcriptional profile specifically in hypothalamic OT neurons of lean but not obese mice

To identify CCK-inducible gene expression modules in a population-specific manner, we explored the effects of peripheral CCK on gene regulatory networks within OT neurons. We generated *OT:RiboTag* mice (OT-*ires-*Cre mice intercrossed to floxed Rpl22^HA^ mice), allowing us to employ translating ribosome <u>a</u>ffinity <u>p</u>urification (TRAP) and RNA sequencing (RNA-seq) of actively translated mRNA selectively in OT neurons (Figure 2D). HA-tagged ribosomes derived from OT neurons were immunoprecipitated (IP) from whole hypothalamic lysate (input) using an anti-HA antibody resulting in an > 80-fold enrichment of *Ot* mRNA in the IP as compared to the input fraction (Figure 2E). Notably, the IP fraction was significantly enriched for *Cckar* mRNA while being de-enriched for *Cckbr* mRNA as compared to the input (Figure 2F). Importantly, mice chronically fed a HFHS diet tended to have lower *Cckar* mRNA and higher *Cckbr* mRNA abundance compared to lean, SC diet-fed control mice (Figure 2G). We next explored more broadly how HFHS diet exposure affects gene expression networks and pathways in OT neurons. Thus, we performed GO enrichment analysis of a total of 3127 differentially expressed genes (DEG), regarded significant with FDR adjusted p-value < 0.05 using the Wald significance test, between vehicle-treated SC diet-versus HFHS diet-fed mice. By employing the cellular component category analysis, we found that HFHS diet-exposure strongly modifies pathways involved in organelle localization, Golgi apparatus function, and synaptic vesicle transport and recycling as well as exocytosis (Figure S2D). Lastly, we injected SC diet- or HFHS diet-fed cohorts of *OT:RiboTag* mice with either CCK (20 μg/kg BW; *i*.*p*.) or vehicle in order to conduct TRAP followed by high-throughput RNA-seq for both input and IP samples. Strikingly, in mice maintained on SC diet, administration of CCK two-hours before sacrifice elicited widespread and profound changes in the gene expression profile of OT neurons compared to vehicle (158 DEG with p-value <0.05 after FDR correction, employing Wald significance test; Figure 2H, I). Of note, the respective input samples did not exhibit any significant gene expression signatures upon CCK delivery further supporting the notion that OT neurons represent a major CCK-responsive population within the hypothalamus (Figure S2E, F). Strikingly, all CCK-related transcriptomic changes were completely abolished if mice were chronically fed a HFHS diet (Figure 2H). In sum, an acute increase of circulating CCK elicits characteristic changes in mRNA translational activity at the level of OT neurons in lean but not in obese mice.

### CCK triggers the activation of PVN^OT^ neurons via CCK_A_R-mediated mechanism in lean but not in obese mice

We next investigated the mechanism of action to discern if PVN^OT^ neurons directly sense CCK levels or whether the response is mediated via relay pathways downstream of a vagal mechanism (Miller et al. 1993; Luckman et al. 1993). Selective expression of a genetically encoded Ca^2+^ indicator, GCaMP6f, was specifically targeted to PVN^OT^ neurons by stereotaxically injecting AAV-DIO-EF1α-GCaMP6f) into OT-*ires*-Cre mice. Using 2-photon excitation Ca^2+^ imaging of ex vivo brain slices, we found that bath application of CCK in the presence of synaptic blockers evoked robust and immediate increases in fluorescent signals in putative magnocellular PVN^OT^ neurons (Figure 3A-D; Figure S3A, S3B; video S1). This indicates that this population is directly responding to CCK with increased cytosolic Ca^2+^ transients. To identify the molecular mechanisms that enable PVN^OT^ neurons to directly sense systemic CCK, we next applied single molecule fluorescent *in-situ* hybridization (FISH; RNAscope) and assessed the expression of the CCK_A_ receptor subtype in PVN^OT^ neurons. CCK_A_R, which is predominantly found in vagal sensory neurons along the alimentary canal (Dourish and Hill 1987; Williams et al. 2016), constitutes the major subtype mediating the food intake-suppressive effect of CCK and has previously been implicated in the activation of OT neurons in rodents (Luckman et al. 1993; Miller et al. 1993). Contrary to the initially proposed vagally mediated mechanism, and consistent with our ribososomal profiling results, we here detected abundant *Cckar* mRNA expression in PVN^OT^ neuronal somata (Figure 3E), which was significantly lower in mice chronically exposed to HFHS diet as compared to SC diet-fed control mice (Figure 3F). To further substantiate a direct CCK_A_R-mediated mechanism in PVN^OT^ neurons, we next conducted whole-cell patch-clamp recordings from putative magnOT neurons *ex vivo*. We identified magnOT cells based on their distinct electrophysiological profile, e.g. characteristic transient outward rectification during depolarizing current injections (Eliava et al. 2016). Superfusion with a potent and selective CCK_A_R agonist (A-71623; 25 nM) elicited robust increases in action potential frequency of putative magnOT neurons from SC diet-fed mice. In contrast, mice chronically fed a HFHS diet did not show any significant increases in action potential frequency when A-71623 nor native CCK-8s (engaging both CCK_A_R and CCK_B_R; 50 nM) were bath applied (Figure 3G, 3H). Notably, magnOT neurons from SC diet-versus HFHS diet-fed mice did not exhibit differences in electrophysiological properties, such as basal action potential frequency, FI curve (firing frequency as a function of injected current), and input resistance (Figure S3C-E). Based on these results, we thus conclude that mice on a SC diet can detect systemic CCK on the level of PVN^OT^ neurons via a direct CCK_A_R-mediated mechanism that becomes aberrant upon feeding mice with a HFHS diet.

**Figure 3:**
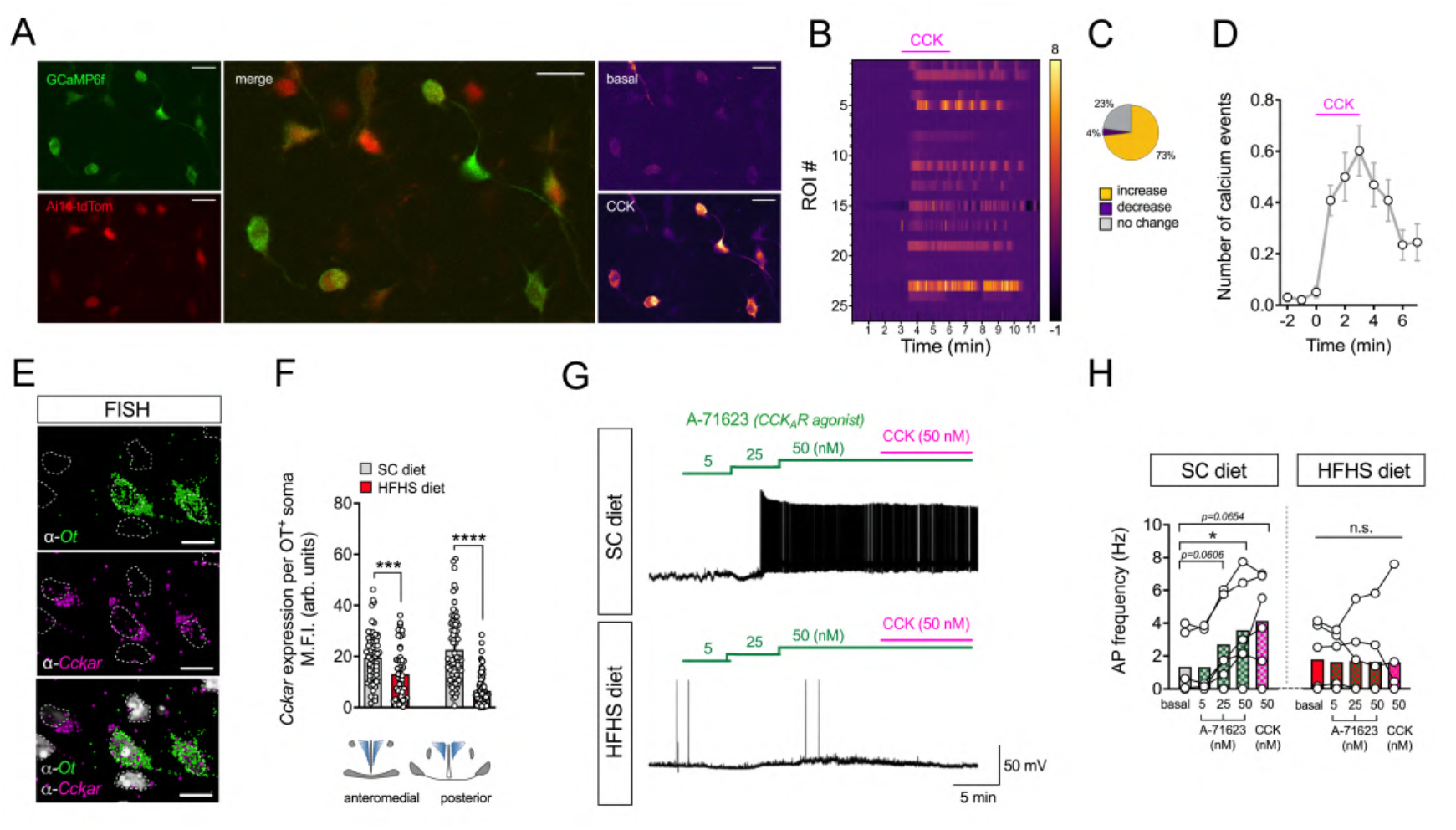
PVN^OT^ neurons are activated by CCK *via* a direct, CCK_A_R-dependent mechanism in lean but not obese mice. **(A)** 2-photon excitation Ca^2+^ imaging of an acute brain slice expressing the genetically-encoded Ca^2+^ indicator GCaMP6f (green; top left panel) in PVN^OT^ neurons conditionally tagged by Ai14-tdTomato (red; bottom left panel) in a cell type-specific manner (merge; middle panel). Representative images taken during the timelapse recording before (basal; top right panel) and after bath-application of 50 nM CCK (CCK; lower right panel) in the presence of synaptic blockers. Scale bars, 25 μm. **(B)** Heatmap representation of cytosolic Ca^2+^ transients of individual PVN^OT^ neurons upon bath-application of CCK (50 nM) in the presence of synaptic blockers. n = 1 mouse, 49 neurons. **(C)** Pie chart diagram illustrating the percentages of PVN^OT^ neurons that increase (yellow) or decrease their activity (purple) upon CCK application, or that do not exhibit any change (gray). **(D)** Quantification of Ca^2+^ events over time displayed in 1-minute bins. n = 1 mouse, 49 neurons **(E)** 3D-rendered, high-power confocal micrograph depicting *Ot* mRNA (green), *Cckar* mRNA (magenta) and DAPI (grey) upon FISH (fluorescent *in situ* hybridization; RNAscope). Individual nuclei are outlined for demarcation within the field-of-view. Scale bar, 20 μm. **(F)** Corresponding quantification of FISH (RNAscope) using background-corrected mean fluorescence intensity (M.F.I.) of *Cckar* mRNA per *Ot*^+^ soma in the rostromedial and caudal PVN of adult male C57BL/6J mice fed either SC or HFHS diet. Data are presented as mean of all somata analyzed ± SEM. **** P < 0.0001, *** P < 0.001. n = 3 mice, 4-8 hemisections per mouse (unpaired Student’s *t*-test). **(G)** Representative traces of action potential frequency of putative magnOT neurons derived from SC diet-fed versus HFHS diet-fed mice in response to increasing concentrations of bath-applied A-71623 (5, 25, and 50 nM) followed by superfusion with native CCK (50 nM). **(H)** Summary of changes in action potential frequency visualized in (G). Data are presented as mean superimposed with individual data points. * P < 0.05, n.s., not significant. n = 2-3 mice/ 6 neurons per mouse (two-way ANOVA).

### Targeted PVN^OT^ neuron activation restores CCK-induced satiety in HFHS diet-fed mice

Based on our previous finding, we hypothesized that CCK’s inability to suppress high-fat diet intake (Covasa and Ritter 1998, 2000; French et al. 1995; Torregrossa and Smith 2003) is a consequence of its decoupling from the hypothalamic OT system, which we have observed upon chronic HFHS diet exposure. To test the pathophysiological relevance of this particular gut-brain pathway, we circumvented this diet-associated impediment of PVN^OT^ neuron engagement by administering systemic CCK to HFHS diet-fed mice while concomitantly activating PVN^OT^ neurons using <u>d</u>esigner receptors <u>e</u>xclusively <u>a</u>ctivated by <u>d</u>esigner <u>d</u>rugs (DREADDs). To accomplish this, we stereotaxically injected either AAV-DIO-hSYN1-hM3Dq-mCherry (an activating DREADD) or a control virus into the PVN of adult male OT-*ires*-Cre mice. After four weeks, mice underwent dosing acclimatisation with sham *i*.*p*. injections within metabolic cages for three consecutive days. Consistent with the literature, CCK administration on the fourth day 10 min before dark onset produced a significant suppression of food intake. In addition, a prolonged feeding latency in SC diet-fed control mice relative to vehicle injection was recorded (Figure 4A); in contrast, CCK injections did not reduce food intake in obese control mice chronically fed a HFHS diet (Figure 4B), as previously observed by others (Covasa and Ritter 1998, 2000; Swartz, Savastano, and Covasa 2010). Next, we asked whether the chemogenetic activation of PVN^OT^ neurons is sufficient to restore CCK-mediated hypophagia in HFHS diet-fed mice that selectively express hM3Dq (hM3Dq^OT+/PVN^ mice). To do this, we pre-injected HFHS diet-fed hM3Dq^OT+/PVN^ mice with the DREADD-activating ligand clozapine-N-oxide (CNO) 20 minutes prior to the systemic administration of CCK and closely assessed their feeding behavior. Strikingly, the simultaneous chemogenetic activation of PVN^OT^ neurons readily restored the sensitivity to CCK and produced a durable suppression of HFHS diet intake over the course of 2 h (Figure 4C). Importantly, in the same cohort of mice, chemogenetic activation of PVN^OT^ neurons without CCK administration did not alter HFHS diet intake relative to control mice (control mice: 1.03 ± 0.19 kcal versus hM3Dq^OT+/PVN^ mice: 0.79 ± 0.17 kcal over 2 hours post CNO). Based on these findings, we inferred that activation of PVN^OT^ neurons is necessary to mediate CCK-induced satiation, but that their activation is not sufficient to suppress HFHS diet intake on its own. To determine how PVN^OT^ neurons are required for co-executing intake suppression with CCK, we next assessed changes in neuronal activity patterns in control mice versus hM3Dq^OT+/PVN^ mice receiving either CNO alone or CNO+CCK (Figure 4D). In control mice, HFHS diet feeding greatly blunted the induction of c-fos immunoreactivity in virus-targeted OT^mCherry+^ neurons by CNO+CCK co-administration relative to CNO alone. In contrast, hM3Dq^OT+/PVN^ mice had robust and near-complete activation of virus-targeted OT^hM3Dq-mCherry^ neurons, regardless if given CNO-only or CNO+CCK in combination (Figure 4E). Importantly, combined CNO+CCK administration in these mice also resulted in a greatly potentiated activation of the PVN overall, including neighboring non-OT neurons (Figure 4F). Thus, this pronounced c-fos response in the PVN upon CNO+CCK, which was observed in hM3Dq^OT+/PVN^ mice but not in control mice or hM3Dq^OT+/PVN^ mice given CNO only, precisely reflected the changes in feeding behavior and further suggests that PVN^OT^ engagement is a prerequisite for broader PVN activation and suppression of food intake upon CCK.

**Figure 4:**
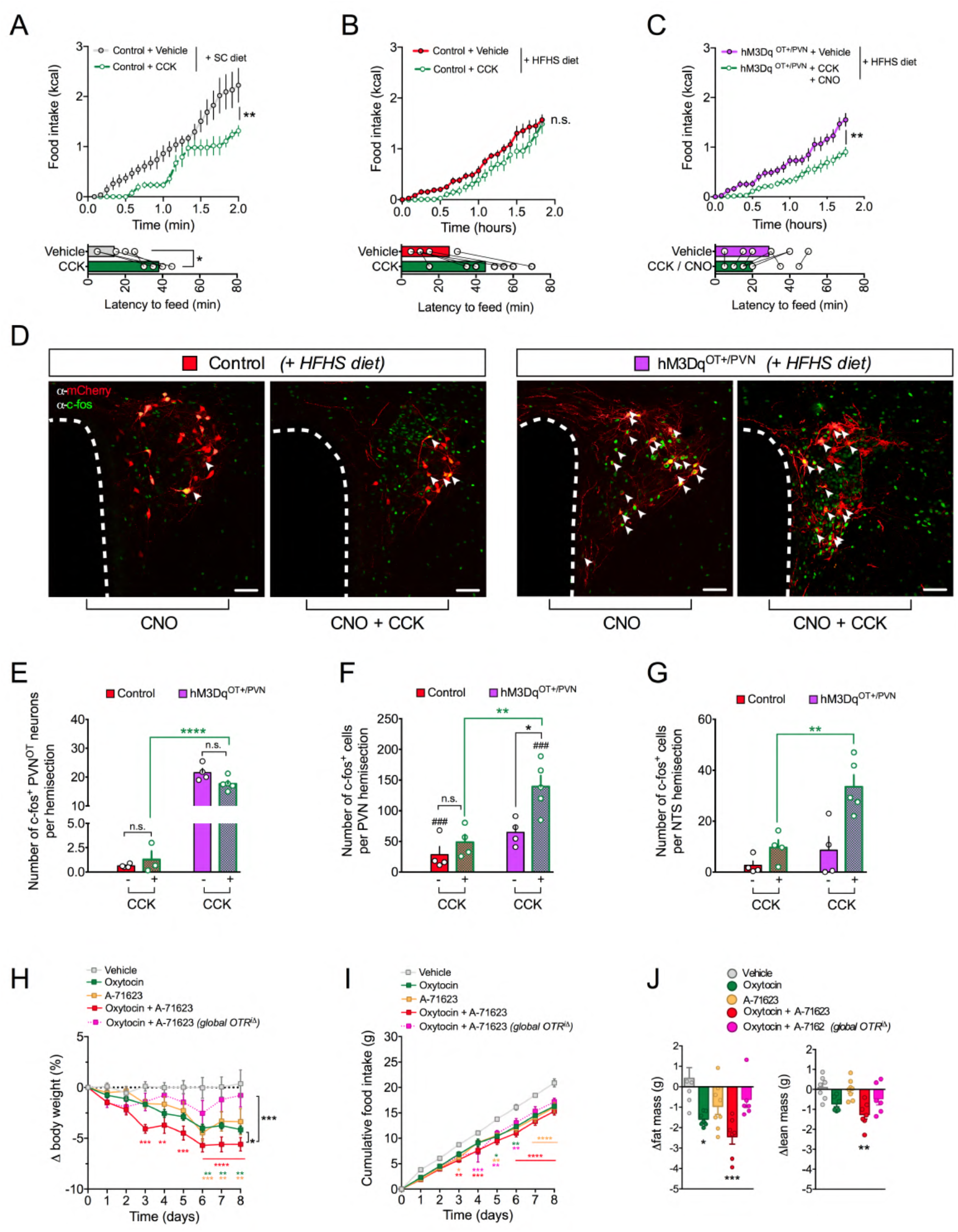
Blunted suppression of food intake in response to CCK on a HFHS diet is reinstated by concomitant chemogenetic activation of PVN^OT^ neurons. **(A)** Cumulative food intake of SC diet-fed control mice (nanoinjected with AAV-hSyn-DIO-mCherry) upon vehicle versus CCK (20 μg/kg BW *i*.*p*.) plus CNO (1 mg /kg BW *i*.*p*.). Data are presented as mean ± SEM. * P < 0.05. n = 5 mice in a cross-over design (two-way ANOVA). **(B)** Cumulative food intake of HFHS diet-fed control mice (nanoinjected with AAV-hSyn-DIO-mCherry) upon vehicle versus CCK (20 μg/kg BW *i*.*p*.) plus CNO (1 mg /kg BW *i*.*p*.). Data are presented as mean ± SEM. n.s., not significant. n = 5 mice in a cross-over design (two-way ANOVA). **(C)** Cumulative food intake of HFHS diet-fed hM3Dq^OT+/PVN^ mice (nanoinjected with AAV-hSyn-DIO-hM3Dq-mCherry) upon vehicle versus CCK (20 μg/kg BW *i*.*p*.) plus CNO (1 mg /kg BW *i*.*p*.). Data are presented as mean ± SEM. * P < 0.05. n = 5 mice in a cross-over design (two-way ANOVA). **(D)** Representative confocal micrographs depicting neuronal activation by means of nuclear c-fos immunoreactivity (green) in virally transduced PVN^OT^ neurons (mCherry^+^; red). Scale bar, 50 μm. **(E)** Quantification of activated (c-fos^+^) virally transduced PVN^OT^ neurons (mCherry^+^) upon CCK (20 μg/kg BW *i*.*p*.) plus CNO (1 mg /kg BW *i*.*p*.) injections in HFHS diet-fed hM3Dq^OT+/PVN^ mice or control mice. n = 4 mice / 3-8 hemisections. **(F)** Quantification of activated (c-fos^+^) PVN neurons overall upon CCK (20 μg/kg BW *i*.*p*.) plus CNO (1 mg /kg BW *i*.*p*.) injections in HFHS diet-fed hM3Dq^OT+/PVN^ mice or control mice. n = 4 mice / 3-8 hemisections. **(G)** Quantification of activated (c-fos^+^) NTS neurons upon CCK (20 μg/kg BW *i*.*p*.) plus CNO (1 mg /kg BW *i*.*p*.) injections in HFHS diet-fed hM3Dq^OT+/PVN^ mice or control mice. n = 4 mice / 3-8 hemisections. **(H)** Relative body weight changes of adult male wildtype mice and tamoxifen-inducible global OTR^-/-^ mice all fed HFHS diet and treated bi-daily with either OT (500 nmol/kg BW; *s*.*c*.), A-71623 (30 nmol/kg BW *i*.*p*.) or their combination. Data are presented in percent of initial body weight as mean ± SEM. * P < 0.05, ** P < 0.01, *** P < 0.001, **** P < 0.0001, n.s. = not significant. n = 6-8 mice (two-way ANOVA). **(I**) Cumulative food intake of adult male wildtype mice and tamoxifen-inducible global OTR^-/-^ mice all fed HFHS diet and treated bi-daily with either OT (500 nmol/kg BW *s*.*c*.), A-71623 (30 nmol/kg BW *i*.*p*.) or their combination. Data are presented as mean ± SEM. * P < 0.05, ** P < 0.01, *** P < 0.001, **** P < 0.0001, n.s. = not significant. n = 6-8 mice (two-way ANOVA). **(J)** Changes in body composition of the cohort shown in (H, I) at the end of study. Data are presented relative to initial body composition as mean ± SEM ± SEM. ** P < 0.01, **** P < 0.0001. n = 5-7 mice (unpaired Student’s *t*-test).

### Chemogenetic PVN^OT^ neuron stimulation potentiates the CCK-induced activation of hindbrain NTS neuron in HFHS diet-fed mice

Intrigued by this broad PVN activation in CNO+CCK-injected hM3Dq^OT+/PVN^ mice, we next pondered whether this effect would extend beyond the PVN also re-sensitizing other critical circuit nodes of central GI hormone signaling. As extensively described elsewhere (Blevins and Baskin 2010), peripheral CCK co-activates a dispersed neurocircuitry including the PVN as well as the nucleus tractus solitarius (NTS) in the caudal brainstem – two regions that are heavily interconnected and are both critical for the orchestration of the behavioral and physiological responses towards CCK (Rinaman et al. 1993; Ueta et al. 2000). In order to interrogate whether HFHS diet-induced aberrations in PVN^OT^ neurons would impact the structure and function of this hypothalamus-brainstem neurocircuit, we employed dual-color 3D whole-brain imaging (iDISCO; (Renier et al. 2014)) of OT:Ai14 reporter mice in combination with immunostaining against tyrosine hydroxylase (TH). By this, we were able to resolve the descending axonal projections from PVN^OT^ neurons as well as their extensive innervation of subregions throughout the NTS, including the catecholaminergic A2/C2 cell group (TH^+^) (Figure S4A-D; video S2). Next, we analyzed the activation state of NTS neurons, including the A2/C2 cell group, upon CNO+CCK co-injections of the same cohort of hM3Dq^OT+/PVN^ mice and controls as in Figure 4. Strikingly, we observed that hM3Dq^OT+/PVN^ mice exhibited significantly more c-fos^+^ neurons in the NTS upon CNO+CCK co-administration relative to control mice (Figure 4G), which was particularly prominent for the catecholaminergic A2/C2 cell group in the caudal medial portion of the NTS (Figure S4E, F). In sum, our data suggests that defective sensing of gut-born CCK upon chronic HFHS diet exposure is a consequence of a disturbed hypothalamus-brainstem network. Importantly, our data further highlights the hitherto unrecognized fact that PVN^OT^ neurons constitute a key population within the hierarchical structure of this network, which is evidenced by the fact that their selective activation readily re-sensitized mice to the feeding-suppressive effect of systemic CCK. We thus conclude that PVN^OT^ neurons function as a pivotal central hub whose targeted activation can restore the transmission of the gut-derived anorexigenic signal CCK under obesogenic conditions.

### Combined OTR-CCK_A_R co-agonism improves metabolic outcomes in diet-induced obese mice

Since the hypothalamic OT system co-executes food intake suppression together with CCK, a process that is disrupted upon HFHS diet exposure, we hypothesized that the pharmacological combination of synthetic OT and a potent and selective CCK_A_R agonist (A-71263) (Asin et al. 1992) would confer significant metabolic improvements in DIO C57BL/6J mice. As compared to respective mono-agonism, co-treatment with OT (500 nmol/kg BW; *s*.*c*.) and A-71263 (30 nmol/kg BW; *i*.*p*.) twice daily for 10 days resulted in increased body weight loss (Figure 4H) which was associated with a significant reduction in food intake relative to what occurred in vehicle-treated control mice (Figure 4I). Body composition analyses further revealed that the substantial weight-lowering effect of combined OT and A-71263 was primarily due to a loss in fat mass (Figure 4J). Importantly, combined OT and A-71263 treatment had no effect on body weight or composition in mice that globally lack OT receptor upon tamoxifen-induced Cre-loxP recombination in adulthood (OTR^iΔ^ mice). Thus, we conclude that integrating OTR- and CCK_A_R agonism promotes favorable metabolic effects in DIO mice and might emerge as a promising new combination to be added to the current arsenal of anti-obesity polypharmacies.

### Single-nucleus RNA-seq2 reveals intersectional regulation of hypothalamic OT neurons by CCK_A_R and κ-opioid receptors that is dependent on dietary context

We next sought to explore the transcriptional diversity across hypothalamic OT neurons at the resolution of single cells. To accomplish this we generated mice in which the nuclei of OT neurons are tagged with super-folded (sf)GFP; specifically, we back-crossed OT-*ires*-Cre mice with a reporter mouse line that Cre-dependently expresses the nuclear membrane protein SUN-domain containing protein 1 (SUN1) fused to sfGFP (*OT:Sun1-sfGFP* mice; Figure 5A). Being therefore amenable to fluorescence-activated cell sorting (FACS), we then isolated hypothalamic OT neuronal nuclei from SC diet- or HFHS diet-fed mice and individually sorted them into 384-well plates (Figure 5B). Per mouse brain, we yielded circa 500 sfGFP^+^ nuclei, which is consistent with stereological counting studies of OT neuron numbers (Lewis et al. 2020). To assess the overall impact of chronic HFHS diet-feeding, we performed differential gene expression analyses (123 DEG with adjusted p-values <0.05) comparing the entirety of single nuclei transcriptomes between mice fed SC diet versus HFHS diet. HFHS diet exposure was associated with substantial changes in gene expression throughout the OT neuronal population (Figure 5C). We then queried our snRNA-seq2 data set for the expression of receptors related to hormonal, neuropeptide and low-molecular transmitter signaling implicated in energy homeostasis. We found multiple OT neurons to express varying combinations of metabolism-related receptors, reassuringly including *Cckar*, suggesting substantial convergence in metabolic information processing at the level of distinct OT neuronal subsets (Figure 5D). We then focused on OT neurons exhibiting transcript intersections between *Cckar* and *Oprk1* (κ-opioid receptor 1; KOR), which constitutes an inhibitory G_i_/G_o_-coupled receptor that stands out by its highly abundant expression across OT neurons. Chronic exposure to a HFHS diet increased the proportion of *Cckar*^*+*^ OT neurons co-expressing *Oprk1* mRNA (Figure 5D, E).

**Figure 5:**
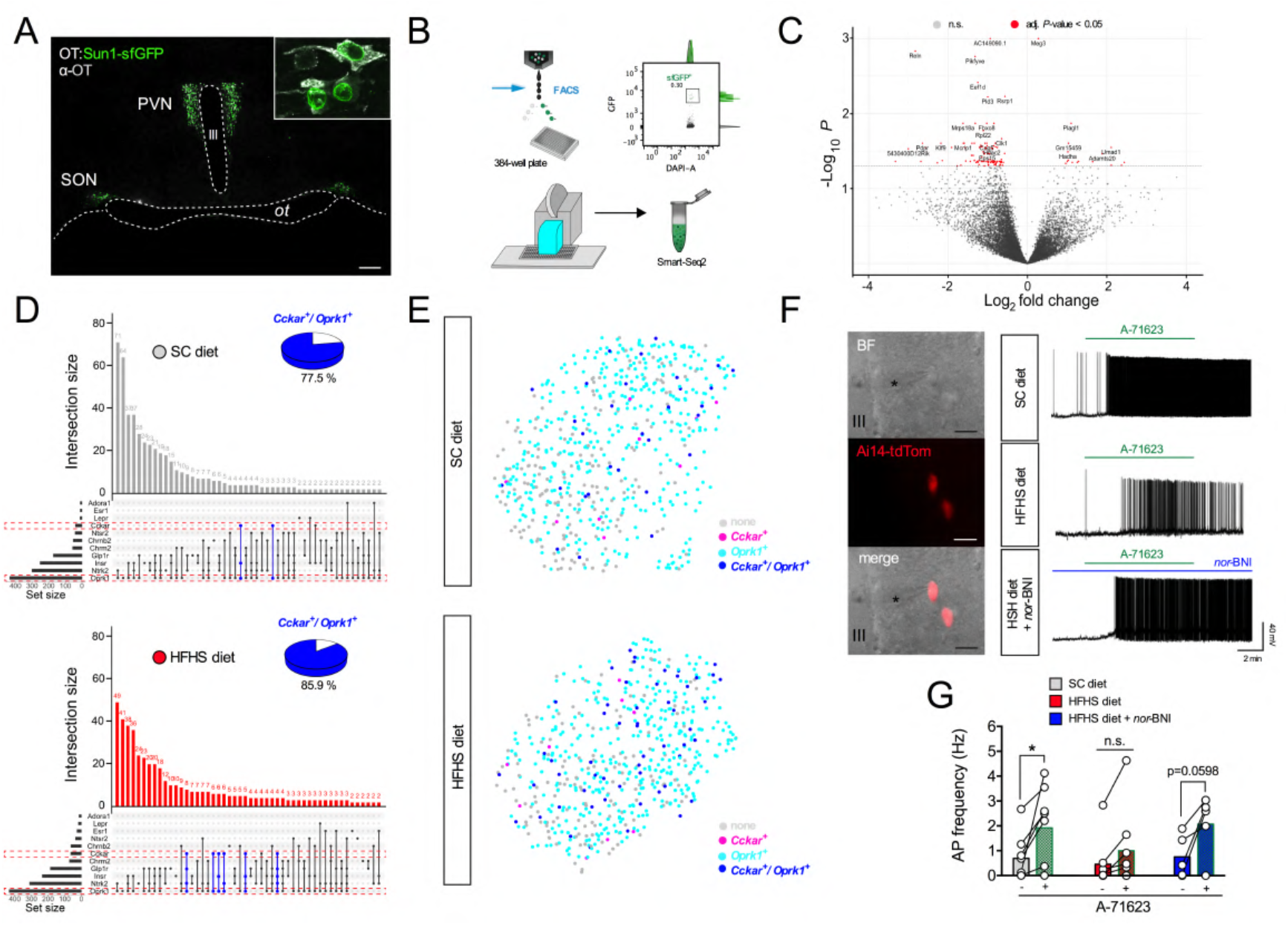
Intersectional regulation of hypothalamic OT neurons by CCK_A_R and κ-opioid receptors is dependent on dietary context. **(A)** Representative confocal micrographs of coronal brain section from an adult male *OT:Sun1-sfGFP* depicting the nuclear localization of sfGFP (green) in hypothalamic OT neurons (grey). Scale bar, 200 μm. **(B)** Schematic illustration of the workflow used to sort individual sfGFP^+^ nuclei into 384-well plates using FACS (top left panel) with representative FACS plot displayed (top right panel); isolation was followed by low-volume pipetting robot-assisted single nuclei lysis, cDNA synthesis and library preparation for snRNA-seq2 (lower panel). **(C)** Volcano plot highlighting differential gene expression changes across the sum of individual OT nuclei from adult male *OT:Sun1-sfGFP* mice chronically fed HFHS diet relative so SC diet-fed littermate controls. n = 2 mice. **(D)** Upse*t* plot visualization of intersectional expression of select receptors in individual OT nuclei from SC diet-fed (upper panel) and HFHS diet-fed (lower panel) *OT:Sun1-sfGFP* mice. Transcripts for *Cckar* and *Oprk1* are highlighted (red dash boxes) as well as their intersections (blue bars). Percentages of *Cckar*^*+*^ OT nuclei also expressing *Oprk1* are visualized as pie charts for each panel. **(E)** UMAP plot visualization of individual OT nuclei from SC diet-fed (upper panel) and HFHS diet-fed (lower panel) *OT:Sun1-sfGFP* mice colored according to their expression of either *Cckar* (magenta), *Oprk1* (cyan), their combination (blue), or none (grey). N = 2; 614 cells (SC diet) and 588 cells (HFHS diet). **(F)** Representative traces of action potential frequency of magnOT neurons derived from adult male *OT:Ai14-tdTomato* reporter mice fed either SC diet or HFHS diet fed mice in response to bath-applied A-71623 (25 nM) with or without pre-treatment with *nor*-BNI (200 nM). Scale bar, 25 μm **(G)** Summary of changes in action potential frequency visualized in (G). Data are presented before and after application of A-71623 as mean ± SEM. * P < 0.05, n.s., not significant. n = 2-3 mice/ 5-8 neurons per mouse (paired Student’s *t*-test).

### κ-opioid tone restrains PVN^OT^ neuronal activation by CCK under HFHS diet feeding

Given that signaling through KOR is associated with inhibitory neuromodulation, we hypothesized that HFHS diet-induced κ-opioid tone in *Cckar*^*+*^ OT neurons might indirectly and tonically blunt their responsivity to systemic CCK. To test this hypothesis, we carried out patch-clamp recordings of PVN^OT^ neurons in acute brain slices derived from mice that selectively express tdTomato in OT neurons (*OT:Ai14-tdTomato*). In agreement with our previous findings, we observed that bath-application of the selective CCK_A_R agonist A-71623 (25 nM) dramatically increased action potential frequency in magnocellular PVN^OT^ neurons from SC diet-fed mice independently of presynaptic inputs (Figure 5F, G and S4A, B), whereas this response was significantly blunted if mice were chronically fed HFHS diet (Figure 5F). However, when brain sections from HFHS diet-fed mice were pre-incubated with the potent and highly selective KOR antagonist *nor*-binaltorphimine (*nor*-BNI; 200 nM in aCSF for 20 min), the CCK_A_R agonist A-71623 was readily able to trigger significant action potentials in PVN^OT^ neurons (Figure 5F, G). Thus, as revealed by our snRNAseq2 data set, these electrophysiological insights further underscored the notion that increased κ-opioid signaling within CCK-sensitive OT subsets underlies their tonically restrained excitability under obesogenic diet.

## Discussion

Gut-to-brain communication is pivotal for the control of whole-body energy homeostasis (Clemmensen et al. 2017) and pharmacological targeting of select pathways within this communication network has recently been found to effectively lower body weight in clinically obese populations (Wilding et al. 2021). Spurred by this translational success story, we sought out to explore alternative brain-targeted gut peptide therapeutics that – if rationally combined – might better mimic the pluri-hormonal physiology of eating (Gribble and O’Rahilly 2021). Here, we have elucidated how the anorexigenic gut peptide CCK fails to suppress food intake under obesogenic diets. Specifically, we observed that feeding a HFHS diet induces aberrations in hypothalamic OT neurons on both the transcriptional and electrophysiological levels by which this pivotal cell population ultimately decouples from gut-borne CCK signaling.

### CCK induces a gene expression signature at the level of PVN^OT^ neurons in lean but not obese mice

By employing affinity purification of tagged ribosomes followed by in-depth translatome profiling, we have identified a highly coordinated translational program of distinct mRNA classes that is induced in hypothalamic OT neurons upon systemic CCK injection. In agreement with what occurs in another population of hypothalamic neurons (Cedernaes et al. 2019), our data add to the evidence suggesting that hormonal cues can readily induce robust and characteristic gene expression signatures in specialized cell types to adjust neuronal physiology to the nutritional state. Besides a pronounced enrichment of mRNA species involved in neuroplasticity including retinoic acid signaling (*Rdh9*) (Shearer et al. 2012), two of the transcripts most robustly induced by CCK were *Exoc3l2 and Trappc9/Nbp*, genes that encode for proteins crucially involved in vesicular trafficking and neuropeptide signaling. Intriguingly, mutations in *Trappc9/Nbp* have previously been linked to a Prader-Willi-like obesity syndrome (PWS) in humans (Marangi et al. 2013; Liang et al. 2020). We further revealed that chronic HFHS diet feeding abolishes the induction of this characteristic gene expression signature, suggesting that long-term caloric excess might uncouple episodic hormonal information from molecular processes promoting OT neuronal plasticity and cell-cell communication. However, future investigations are yet required to reveal further intricacies as to how hormone-inducible gene expression signatures modulate hypothalamic function and energy homeostasis and whether they might bear therapeutic potential for dietary and/or genetic forms of obesity.

### Targeted co-activation of OT and CCK signaling elicits enhanced food intake suppression and weight loss despite obesogenic diet exposure

Physiological energy homeostasis crucially depends on the gastrointestinal system informing the brain about the nutritional status quo in the form of gut-derived humoral and neural signals such as CCK (Clemmensen et al. 2017). CCK, as the first-described and paradigmatic anorexigenic gut hormone, suppresses feeding in a variety of species ranging from rodents (Gibbs, Young, and Smith 1973; Crawley et al. 1981) to primates (Falasco, Smith, and Gibbs 1979) including humans (Sturdevant and Goetz 1976). However, its hoped-for therapeutic potential as an anti-obesity drug was questioned based on early observations reporting that chronic intake of fat-rich diets greatly blunts its anorexigenic effect (Covasa and Ritter 1998, 2000; French et al. 1995; Torregrossa and Smith 2003; Beutler et al. 2020). Despite chronic exposure to an obesogenic diet, mice administered CCK had fully restored food intake suppression due to artificially reinstated OT signaling using chemogenetic or polypharmacological means. Besides PVN^OT^ neurons, prior studies have suggested that the nucleus tractus solitarius (NTS) in the brainstem constitutes another pivotal brain region that mediates CCK-induced hypophagia (Brown et al. 1998; Ho et al. 2014; Rinaman and Rothe 2002; Olson et al. 1992). Within the NTS, the catecholaminergic A2/C2 cell group in particular is robustly activated by systemic CCK injections (Rinaman et al. 1993). Notably, this group of brainstem cells is reciprocally interconnected with the hypothalamic OT system and receives dense innervation from parvOT neurons residing in the PVN (Blevins et al. 2003). Conversely, ascending catecholaminergic projections emanating from the NTS target the PVN. In fact, norepinephrine release within the hypothalamus is highly correlated with systemic OT secretion following peripheral CCK injections (Brown et al. 1998), strongly implying that PVN^OT^ neurons and the A2/C2 cell group constitute a pivotal, interrelated tandem for processing CCK-mediated information. Besides the catecholaminergic A2/C2 cell group, the NTS additionally harbors neurons that produce glucagon-like peptide 1 (GLP-1), another potent anorexigen, whose central signaling profoundly intersects with both OT (Rinaman and Rothe 2002; Brierley et al. 2021) and CCK pathways (Borgmann et al. 2021). Together, these novel insights into the intricate crosstalk among various gut-to-brain-pathways make it ever clearer that we are still far from a comprehensive understanding of the poly-hormonal control of eating. Notably, our data now add to the body of literature whose emergent complexity demands a deeper understanding of how various gut peptide signaling axes intersect at the central level and how these interactions are altered in disease states such as obesity.

### Obesity is associated with increased *Oprk1* mRNA expression and inhibitory κ-opioid signaling in PVN^OT^ neurons that restrains their excitability in response to CCK

By devising a novel approach for single-nuclei isolation in combination with snRNA-seq2, we mapped the molecular heterogeneity across the entire hypothalamic OT system. We demonstrated that multiple individual OT neurons express a series of different metabolism-related receptors, suggesting distinct populations with partly overlapping sensitivities towards metabolic cues. One of the most abundant receptors, *Oprk1*, which encodes for the κ-opioid receptor type 1 that is typically associated with inhibitory neuromodulation and is increased in *Cckar*^*+*^ OT neurons by chronic HFHS diet exposure, was highly expressed. *Oprk1* mRNA expression additionally overlapped pronouncedly with that of *Fam19a1* (family with sequence similarity 19, member 1a), a brain-enriched and metabolically responsive neurokine (Lei et al. 2019) that has been associated with human obesity and insulin resistance in a recent genome-wide association study (Rausch et al. 2018). Prompted by the striking *Oprk1* expression levels as well as by the suggested crosstalk with CCK_A_R signaling, we went on to functionally probe for tonic inhibition of CCK-evoked PVN^OT^ neuron activation under HFHS feeding using *ex-vivo* electrophysiology. Indeed, we were able to restore excitability towards CCK_A_R agonism by selectively blocking KOR signaling in brain slices of HFHS diet-fed mice. Consistent with previous reports (Onaka et al. 1995; Leng, Dye, and Bicknell 1997), we now provide additional evidence for the functional significance of increased κ-opioid tone onto hypothalamic OT neurons and propose that it is a major mechanism that contextually restrains the CCK-evoked activation under obesogenic diets. The elucidation of this hitherto elusive mechanism might shed new light on prior observations that pharmacological blockade of opioid signaling specifically in the PVN reduces both homeostatic and hedonic feeding in rats (Koch et al. 1995) and on the fact that mice globally lacking KOR (Oprk1^-/-^ mice) are protected from developing dietary obesity (Czyzyk et al. 2010). Further *in-vivo* studies will be required to disentangle the functional intersections between the OT system, CCK, endogenous opioids, and perhaps other factors such as FAM19A1, in the context of metabolic disorders. In sum, we have shed new light on the molecular heterogeneity within the OT system as well as on its (mal)adaptive plasticity that occurs under (patho)physiological contexts. Future studies will be required to ascertain if these insights might aid to harness OT as such, or in combination with synergistic hormones like CCK and/or KOR blockers, for the development of next-generation precision medicine to correct metabolic physiology.

## Conclusion

In conclusion, we have identified the hitherto elusive mechanisms as to why systemic CCK fails to activate hypothalamic OT neurons under obesogenic diets and delineated approaches such as endogenous opioid modulation to recouple CCK signaling with hypothalamic OT pathways for restoration of gut-brain satiation signaling. These experiments ultimately reinforce the concept that PVN^OT^ neurons constitute a molecularly and functionally diverse assembly of neurons that are tightly enmeshed within a widespread metabolic control network in which they occupy a key position completing a feedback loop between afferent peripheral signals, central circuits and behavior.

## Acknowledgments

The authors thank Cassie Holleman, Elisavet Lola, Clarita Layritz and Nicole Klas for their excellent technical assistance. We thank Dr. Sandrine Lefort for contributing to Ca^2+^ imaging and Maria Richter for critical feedback regarding bioinformatic analyses. This work was supported in part by funding to R.E.C. and P.T.P. from Marie Skłodowska-Curie Grant (ChroMe # 675610), I.G.-G. is a recipient of a fellowship from European Union’s Horizon 2020 research and innovation program under the Marie Sklodowska-Curie actions (842080 – H2020-MSCA-IF-2018), R.H.W received funding from the European 748 Research Council ERC under the European Union’s Horizon 2020 research and innovation 749 programme (#715933), C.P.M.J. and E.S.Q. were supported by the Helmholtz Pioneer Campus, V.G. received funding from German Research Foundation DFG (GR 3619/13-1, GR 3619/15-1, GR 3619/16-1, DFG TRG (GRK) 2174, and SFG Consortium 1158-2), and M.H.T. and C.G.-C. received funding from the European Research Council ERC (AdG grant Hypoflam # 695054 and STG grant AstroNeuroCrosstalk # 757393 respectively), and the German Research Foundation DFG under Germany’s Excellence Strategy within the framework of the Munich Cluster for Systems Neurology (EXC 2145 SyNergy – ID 390857198) and Helmholtz Excellence Network. T.G. and C.G.-C. received funding from the German Center for Diabetes Research (DZD) twinning grant 2020.

## Author Contributions

T.G., C.G.-C., and V.G. conceptualized all studies and designed all experiments. T.G. (surgeries, immunohistochemistry, imaging, qPCR), T.G., F.L., I.G.-G. and O.L.T. (metabolic phenotyping), R.E.C. (FACsorting), E.S.Q. (library preparation snRNA-seq2), V.M. (bioinformatics), and C.D.B.M. (electrophysiology), conducted the experiments, collected, and analyzed the data. T.G., C.G.-C., and V.G. wrote the manuscript in discussion with M.H.T., C.P.M.J., R.H.W., T.D.M., S.C.W., and P.T.P., who revised the article critically for important intellectual content. All authors have read and approved the final version of the manuscript.

## Material and Methods

### Animals

Animal studies were approved by the Animal Ethics Committee of the government of Upper Bavaria (Germany). Wildtype mice (C57BL/6J, Janvier, Le Genest-Saint-Isle, France) or genetically modified mice at adult age (>12 weeks) were provided *ad libitum* access to either a pelleted standard chow (SC) diet (5.6% fat; LM-485, Harlan Teklad) or a high-fat, high-sucrose (HFHS) diet (D12331; 58% of calories from lipids; Research Diets, New Brunswick, NJ). Animals had continuous free access to water and were maintained at 23°C with constant humidity on a 12-h light–dark cycle. OT-*ires*-Cre (also known as *OXT-IRES-Cre*) were originally provided from Jackson Laboratory (strain name: B6;129S-Oxt^tm1.1(cre)Dolsn/^J; # 024234); homozygous OT-*ires*-Cre males were interbred with non-Cre-carrying female mice to obtain experimental cohorts of male mice containing the knock-in allele in heterozygosity. RiboTag mice (strain name: B6N.129-Rpl22^tm1.1Psam^/J; # 011029), Ai14 (strain name: B6.Cg-Gt(ROSA)26Sor^tm14(CAG-tdTomato)Hze^/J; # 007914), ROSA-DTA (strain name: B6.129P2-Gt(ROSA)26Sor^tm1(DTA)Lky^/J; # 009669), and CAG-Sun1sfGFP (strain name: B6;129-Gt(ROSA)26Sor^tm5(CAG-Sun1/sfGFP)Nat^/J; # 021039) were all provided from Jackson Laboratory and used in heterozygosity in final cohorts.

### Genotyping of mouse lines

Ear tags were obtained from mice at the age of 3 weeks and DNA was isolated by boiling the eartags for 30 min in 200 μl 50 mM NaOH at 95°C (ThermoMixer C, Eppendorf). Afterwards, 20 μl 1 M Tris was added to normalize the pH. 2 μl of isolated genomic DNA was used for the genotyping PCR (Promega) using respective protocols.

### Physiological measures and metabolic phenotyping

For assessing acute feeding behavior, mice were individually housed in metabolic cages (TSE PhenoMaster; TSE systems). After 48 h acclimatization, food was removed daily for 3 hours (3pm-6pm) and mice received sham injections with Vehicle (0.9% NaCl) for two consecutive days. On day three, all mice received first CNO (1 mg/kg BW; *i*.*p*.; 25 minutes before dark onset) and CCK-8s (20 μg/kg BW; *i*.*p*.; 10 minutes before dark onset). With dark onset, food hoppers were given back and food intake was automatically measured every 5 minutes. Normal baseline food intake (“Vehicle”) is represented as the calculated mean intake upon the two sham injections. Mice that displayed food spilling were excluded from analysis. Cumulative long-term food intake was measured by co-housing mice (two per cage depending on cohort size). Body composition (fat and lean mass) was assessed by using a magnetic resonance whole-body composition analyzer (Echo-MRI, Houston, TX). Energy expenditure and respiratory exchange ratio (RER) of individual mice were analyzed within metabolic cages of a combined indirect calorimetry system (TSE PhenoMaster; TSE systems). Food and water intake, O_2_ consumption, CO_2_ production, and locomotor activity (*i*.*e*., horizontal and vertical beam breaks) were measured in 5-minute intervals. EE (kcal/h) was calculated based on the Weir equation (3.815*10^−3^*VO_2_+1.232*10^− 3^*VCO_2_) and total EE (kcal) was correlated to the mean of body weight measured before and after the measurement (Tschop et al. 2011). Glucose tolerance was assessed by the intraperitoneal administration of a glucose bolus (2 g/kg BW; 20% w/v in 1xPBS pH 7.4). Before the glucose tolerance test, mice were fasted for 4 h and glycemia was measured by sampling blood from the tail vein before (0 min) and at 15, 30, 60 and 120 min post injection via a handheld glucometer (Abbott, Wiesbaden, Germany). To assess insulin sensitivity, additional blood was collected at 0 min using EDTA-coated microvette tubes (Sarstedt, Nürnbrecht, Germany) to obtain plasma (5,000 x g for 10 min at 4°C). Insulin concentration was determined using a commercial insulin ELISA following the manufacturer’s instructions (Ultra-sensitive Mouse Insulin ELISA Kit, #90080 Crystalchem, Netherlands). HOMA-IR was calculated using the formula: HOMA-IR=[fasting insulin (mU/l) * fasting glucose (mg/dl)] / 405 (Matthews et al., 1985). Glycated HbA_1C_ was analyzed using DCA Vantage® Analyzer (Siemens, City, Germany).

### *Ex-vivo* brain slices preparation

Adult male mice were sacrificed by cervical dislocation and the brain quickly ablated and placed in ice-cold artificial cerebrospinal fluid (aCSF) modified for slicing, containing (in mM): 87 NaCl, 2.69 KCl, 1.25 NaH_2_PO_4_, 26 NaHCO_3_, 7 MgCl_2_, 0.2 CaCl_2_, 25 D-glucose, and 75 sucrose (330 mOsm/Kg H_2_O, pH 7.4 bubbled with a carbogen mixture of 95% O_2_ and 5% CO_2_). The specimen was glued to a sectioning stage and submerged in ice-cold slicing aCSF in a vibratome (VT1200; Leica Biosystems) chamber. Coronal brain slices (250 μm) containing the paraventricular nucleus of the hypothalamus were sectioned and incubated at 32-33º for 30 min in aCSF, containing (in mM): 124 NaCl, 2.69 KCl, 1.25 NaH_2_PO_4_, 26 NaHCO_3_, 1.2 MgCl_2_, 2 CaCl_2_, 2.5 D-glucose, and 7.5 sucrose (298 mOsm/Kg H_2_O, pH 7.35 constantly bubbled with a carbogen mixture of 95% O_2_ and 5% CO_2_). After this period, the slices were kept under the same conditions but at room temperature for at least 40 min until electrophysiological recordings.

### 2-photon excitation calcium imaging

Cytosolic calcium levels from PVN^OT^ neurons conditionally tagged by Ai14-tdTomato within acute coronal brain slices (250 μm) of mice were monitored by 2-photon excitation microscopy using the genetically encoded calcium indicator GCaMP6f. Single brain slices were transferred to a chamber mounted on a stage of an upright multiphoton laser scanner microscope (FVMPE-RS system, Olympus) and continuously perfused with bubbled-aCSF using a gravity-driven perfusion system at a rate of ∼3 mL/min in the presence of synaptic blockers (20 μM CNQX, 50 μM D-AP5, and 100 μM picrotoxin). Neurons were visualized with a 25x water immersion objective. Excitation illumination was generated by an InSight X3 DUAL tunable laser system (Spectra-Physics). The FluoView image acquisition software (FV31S-SW, Olympus) was used to tune laser emission wavelength to 930 and 1045 nm in order to obtain 2-photon absorption signals from GCaMP6f and tdTomato fluorophores, respectively, at an acquisition rate of 0.5 Hz. Calcium imaging from PVN^OT^ neurons consisted of a 3-min baseline recording followed by bath application of CCK (50 nM) and drug washout. At the end of each experiment, 20 mM KCl was bath-applied to check neuronal viability and calcium signal integrity. Only neurons that responded to KCl were used for analysis. Calcium transients were estimated as changes in GCaMP6f-based fluorescence intensity over the baseline (ΔF/F_0_), considering a calcium event when ΔF/F_0_ > 3 standard deviations greater than the baseline fluorescence signal. The number of calcium events were then plotted over time grouped into 1-min bins in order to quantify changes in the frequency of calcium events in PVN^OT^ neurons in response to CCK application.

### Electrophysiological recordings

Single brain slices were transferred to a chamber mounted on a stage of an upright microscope (BX51WI; Olympus) coupled with a video camera (Prime 95B; Teledyne Photometrics) and continuously perfused with carbogen-bubbled aCSF at a rate of ∼2 mL/min by a gravity-driven perfusion system. Neurons were visualized under infrared differential interference contrast (IR-DIC) optics with a 20x immersion objective (2x post-magnification) using the μManager 1.4 software (Edelstein et al. 2010). Accordingly, Ai14-tdTomato^+^ neurons were identified by epifluorescence-based signals excited at 555 nm wavelength (LedHUB; Omicron-Laserage Laserprodukte GmBH). Patching pipettes were made with thick-walled borosilicate glass (GC150TF-10; Harvard Apparatus) pulled using a horizontal puller (P-97; Sutter Instruments) and filled with an internal solution, containing (in mM): 128 K-gluconate, 8 KCl, 10 HEPES, 0.5 EGTA, 4 Mg-ATP, 0.3 Na-GTP, and 10 Na-phosphocreatine (295 mOsm/Kg H_2_O, pH 7.3), resulting in a pipette tip resistance between 4 and 10 MΩ. Whole-cell recordings were performed with a MultiClamp 700B amplifier (Molecular Devices) in current-clamp mode. Data were acquired at 10-20 kHz and low-pass filtered at 5 kHz (Bessel) with a Digidata 1550B (Molecular Devices) using the Clampex 11.2 acquisition software (pClamp; Molecular Devices). Parvocellular neurons were identified by the presence of a transient outward rectifying current triggered by the application of a hyperpolarizing pulse (ranging from -60 to -40 pA for 1 s) followed by positive current injections (from 20 to 100 pA in five steps of 1 s) (Tang et al. 2020). Membrane potential was monitored in response to bath-applied drugs at I = 0. In some experiments, 6-Cyano-7-nitroquinoxaline-2,3-dione (CNQX; 20 μM), D-(-)-2-Amino-5-phosphonopentanoic acid (D-AP5; 50 μM), and picrotoxin (100 μM) were applied to block fast neurotransmission. Membrane input resistance was measured by the slope of the curve obtained by the response of the membrane potential to injected negative currents (from - 40 to -60 pA in three steps of 1 s). Data were visualized and analyzed using custom-written codes in MATLAB (MathWorks).

### Adeno-associated viruses (AAV) and stereotaxic surgery

In order to assess PVN^OT^ neuronal activity, we conducted stereotaxic surgeries to target the fluorescent Ca^2+^ indicator GCaMP6f to PVN^OT^ neurons in OT-*ires*-Cre mice. AAV2/1-DIO-CAG-GCaMP6f was purchased from Addgene (# 100839, titer: 1.4 × 10^13^ gc/ml). To chemogenetically activate PVN^OT^ neurons by means of DREADD we employed AAV2/1-DIO-hSYN1-hM3Dq-mCherry (Addgene # 44361; titer: 1.6 × 10^13^ gc/ml) versus a respective control virus AAV2/1-DIO-hSYN1-mCherry (Addgene # 50459; titer: 9 × 10^12^ gc/ml). To ablate PVN^OT^ neurons, we produced AAV2/1-OTp-*i*Cre (titer: 2.3 × 10^12^ gc/ml) and AAV2/1-OTp-Venus (titer: 1.2 × 10^13^ gc/ml) according to our previously published protocol (Knobloch et al. 2012), which contain a synthetic 2.6 kb OT promoter element (OTp) faithfully restricting viral transgene expression to OT neurons. Respective AAVs were injected bilaterally (300 nl per hemisphere) using a stereotaxic system combined with a binocular 3.5x-90x stereomicroscope (AMScope, USA). Mouse skull was exposed via a small incision and a craniotomy above the viral injection sites was performed using a micro-precision drill. A pulled glass pipette with a 20-40 μm tip diameter was carefully lowered to reach the PVN on each side of the brain, respectively, and AAVs were applied at low speed using manually-applied air pressure. Undesired diffusion of viral particles was further prevented by slow retraction of the glass pipette 5 min post injection. The following stereotaxic coordinates were obtained from ‘The Mouse Brain in Stereotaxic Coordinates’ (Franklin and Paxinos, 2019) and used to target the PVN of the mouse brain: −0.7 mm posterior and ±0.2 mm lateral to the bregma and −4.75 mm ventral from the dura mater. Anesthesia was performed by a mixture of ketamine and xylazine (100 mg/kg body weight and 7 mg/kg bodyweight, respectively) while acute Metamizol (200 mg/kg, subcutaneous) followed by Meloxicam (1 mg/kg, on three consecutive days, subcutaneous) was administered for postoperative analgesia.

### Transcardial perfusion, brain sectioning and immunohistochemistry

Animals were sacrificed with CO_2_ and transcardially perfused with 20 ml phosphate-buffered saline (PBS) (GibcoTM, pH 7.4) by using a peristaltic pump at 120 mmHG (Instech, High Flow P720 equipped with 21G canula). Perfusions were finalized with 20 ml of 4% paraformaldehyde (PFA) in PBS, pH 7.4, brains were removed and post-fixed in 4% PFA at 4°C overnight. The following day, brains were then equilibrated with 30% sucrose in Tris-buffered saline (TBS, pH 7.2) for 48 h before being sectioned into 40 μm coronal slices using a cryostat (CM3050S; Leica, Germany). Three to four brain sections per mouse were selected containing the middle portion of the PVN or NTS and further subjected to additional immunohistochemistry. Brain sections were first washed with TBS and incubated overnight at 4°C with primary antibodies in a solution containing 0.25% porcine gelatine and 0.5% Triton X-100 in TBS, pH 7.2. The next morning, sections were serially rinsed in TBS, pH 7.2, and incubated with respective secondary antibodies diluted in TBS, pH 7.2 containing 0.25% porcine gelatine and 0.5% Triton X-100 for 2h. Sections were serially washed in TBS with the last washing additionally containing fluorescent Nissl stain (1:100 NeuroTraceTM 435/455; ThermoFisher, Germany) in order to identify neuronal cells and anatomical demarcations. Lastly, brain sections were mounted on gelatine-coated glass slides, dried and cover-slipped with a polyvinyl alcohol (mowiol®, Merck, Germany) mounting medium supplemented with DABCO (Merck, Germany).

### Fluorogold injections

To distinguish magnOT and parvOT neurons, fluorogold (15 mg/kg BW; *i*.*p*.) was administered to adult male *OT:Ai14* reporter mice maintained on SC diet. Mice were then perfused 7 days post injection of fluorogold following three hours of food removal plus two additional hours after CCK-8s injection (20 μg/kg BW; *i*.*p*.).

### CCK-8s injections

For all *in-vivo* experiments, food was removed for 3 hours before dark onset (3– 6 pm) in order to normalize feeding behavior. Cholecystokinin-octapeptide, sulfated (CCK-8s; Tocris, St. Louis, USA) was reconstituted in 0.9% NaCl and administered 10 min before dark onset (20 μg/kg BW; *i*.*p*.).

### iDISCO-based whole brain clearing

Clearing protocol was adopted from (Renier et al. 2014)) with slight adjustments. In brief, *OT:Ai14* mice were transcardially perfused with 1xPBS (pH 7.4) followed by 4% PFA and their brains were carefully removed. Following post-fixation in 4% PFA overnight at 4°C, brains were washed three times in 1 x PBS (0.2 % TritonX-100; v/v) for 1 h at room temperature. Brains were pre-treated by incubation in 1 x PBS (0.2 % TritonX-100 and 20 % DMSO; v/v) for 48 h shaking at 37°C followed by 1 x PBS (0.1 % Tween-20, 0.1 % TritonX-100, 0.1 % Deoxycholate, 0.1 % NP40 and 20 % DMSO; v/v and w/v, respectively) for 48 h shaking at 37°C. After washing brains three times in 1 x PBS (0.2 % TritonX-100; v/v) for 1 h at 37°C, they were incubated in permeablization solution (1 x PBS with 0.2 % TritonX-100, 0.3 mM glycine, and 20 % DMSO; v/v and w/v, respectively) for 48 h shaking at 37°C. Thereafter, brains were incubated in blocking solution (1 x PBS with 0.2 % TritonX-100, 3 % donkey serum, 3 % rabbit serum, and 10 % DMSO; v/v) for 48 h shaking at 37°C. After washing brains four to five times in 1 x PBS (0.2 % TritonX-100, and 10 μg/ml heparin; v/v) they were incubated with the primary antibody (rabbit-anti-dsRed at 1/300 dilution) in 1 x PBS with 0.2 % TritonX-100, 10 μg/ml heparin, 3 % rabbit serum, and 5 % DMSO (v/v) for four days shaking at 37°C. After washing brains four to five times in 1 x PBS (0.2 % TritonX-100, and 10 ug/ml heparin; v/v) they were incubated with the secondary antibody (donkey-anti-rabbit Alexa Fluor 647) in 1 x PBS with 0.2 % TritonX-100, 10 μg/ml heparin, 3 % donkey serum, and 5 % DMSO (v/v) for four days shaking at 37°C. After immunolabeling, brains were washed again for four to five times in 1 x PBS (0.2 % TritonX-100, and 10 μg/ml heparin; v/v). Brains were cleared by an ascending dilution of methanol/H_2_O (20 %, 40 %, 60 %, 80 %, 100 %) for 1 hour each at room temperature and were left in fresh 100 % methanol overnight. Delipidation was achieved by incubation in 66 % dichloromethane (DCM) and 33 % methanol for 6 hours at room temperature. After short incubations of brains in 100 % DCM, they were finally placed in the refractive index matching solution consisting of 100 % dibenzylether (DBE) until imaging.

### Translating ribosome affinity purification (TRAP)

Adult male *OT:RiboTag* mice with both transgene alleles in heterozygosity were fed either SC diet or HFHS diet for 3 months. On the day of experiments, food was removed for 3 hours (3 pm – 6 pm) and mice received either vehicle or CCK-8s (20 μg/kg BW; *i*.*p*.; Tocris, St. Louis, USA) injection at dark onset (6 pm) and were sacrificed 2 hours post injection. Hypothalami were rapidly removed, snap-frozen and stored at -80°C until further processing. Per sample, two hypothalami were combined and processed according to previously published protocol (Sanz et al. 2009). Yield of input and immunoprecipitate were independently quantified using a Bioanalyzer (Agilent RNA 6000 Pico Kit; Agilent Technologies, Santa Clara, USA) and Quant-IT RiboGreen Kit (ThermoFisher Scientific Inc., Rockford, IL USA) according to the manufacturer’s instructions. Samples were amplified and synthesized into cDNA according to the manufacturer’s protocol using the SMART-Seq® v4 Ultra® Low Input RNA Kit for Sequencing (Takara Bio Inc., Shiga, Japan) and amplification yield was checked using a Bioanalyzer (Agilent High Sensitivity DNA Kit, Agilent Technologies, Santa Clara, USA). Single-indexed libraries were generated using ThruPLEX® DNA-seq Kit (Takara Bio Inc., Shiga, Japan), pooled and checked again using a Bioanalyzer (Agilent High Sensitivity DNA Kit, Agilent Technologies, Santa Clara, USA).

### TRAP transcriptomics analyses

Sequencing was performed at the Helmholtz Zentrum München (HMGU) by the NGS-Core Facility. After a final quality control, the libraries were sequenced in a paired-end mode (2×150 bases) in the Novaseq6000 sequencer (Illumina) with a depth of ≥ 40 Million paired reads per sample. BCL files were converted to FASTQ using Bcl2fastq v.2.20. The alignment was done using STAR v.7.2a and was mapped to the mouse reference genome GRCm38 (mm10). The counts and FPKM were generated using HTSeq v0.11.2 For transcriptomics downstream analysis, sample distance matrix was generated using pheatmap package from R (Kolde 2015). Differential expression analysis results were illustrated in the form of volcano plots using R-package EnhancedVolcano (Blighe, Rana, and Lewis 2021). The expression profile of DEGs over the samples was shown in the heatmap using R-package pheatmap (Kolde 2015). GO enrichment analyses were performed, and the results were illustrated using clusterProfiler (Yu et al. 2012) and ggplot2 (Wickham 2016) R-packages, respectively.

### RNA isolation and qPCR analysis

RNA was isolated from tissues using a commercially available kit (MicroRNeasy Kit, Qiagen, Hilden, Germany). Identical amounts of RNA were reverse-transcribed to cDNA using Superscript III (Invitrogen, Darmstadt, Germany) and gene expression was analyzed using TaqMan probes (ThermoFisher Scientific Inc., Rockford, IL USA) using a ViiATM7 Real Time PCR System or QuantStudio 6 FLEX Real Time PCR System (ThermoFisher Scientific Inc., Rockford, IL USA). Expression changes were calculated using the 2^-ΔΔCt^ method normalized by *Hprt* as housekeeping gene. When indicated, qPCR expression analysis was conducted on cDNA derived from immunoprecipitated RNA of *OT:RiboTag* mice following reverse (SMARTer® PCR cDNA synthesis kit; Takara Bio Inc., Shiga, Japan).

### Isolation of nuclei tagged in specific cell types (INTACT)

CAG-Sun1-sfGFP mice (INTACT mice) were crossed with OT-*ires*-Cre mice to generate heterozygous mice (Deal and Henikoff 2011). Whole hypothalami were individually processed to obtain single nuclei following a previously described protocol (Krishnaswami et al. 2016) with slight modifications. Briefly, frozen hypothalami were transferred to a Dounce homogenizer containing 1mL of freshly prepared ice-cold nuclei isolation buffer (0.25M sucrose, 25 mM KCl, 5 mM MgCl_2_, 20mM Tris pH 8.0, 0.4% IGEPAL 630, 1 mM DTT, 0.15 mM spermine, 0.5 mM spermidine, 1x phosphatase & protease inhibitors, 0.4 units RNasin Plus RNase Inhibitor, 0.2 units SuperAsin RNase inhibitor). Homogenization was achieved by carefully douncing 10 strokes with the loose pestle, incubating on ice for 5 min and further douncing 15 more strokes with the tight pestle. The homogenate was filtered through a 20 μm cell strainer, centrifuged at 1000 x g for 10 min at 4°C, the nuclei pellet resuspended in 450 μl of staining buffer (PBS, 0.15 mM spermine, 0.5 mM spermidine, 0.4 units RNasin Plus RNase Inhibitor, 0.4% IGEPAL-630, 0.5% BSA) and incubated for 15min on ice. Nuclei pellets were resuspended in 1 mL of fresh staining buffer supplemented with DAPI 1 μg/μL. Nuclei integrity was assessed in the DAPI channel under a Zeiss microscope (Axio Scope, Zeiss, Germany). Doublet discrimination and DAPI staining were used for appropriate gating of single nuclei and the signal on the 488 (FITC) channel of the IgG-isotype control determined the adequate gating of GFP^-^ and GFP^+^ nuclei.

### Fluorescence-assisted cell sorting

GFP^+^ nuclei were sorted with a 70 μm nozzle into 384-well PCR plates (thin-walled, BioRad, HSP3901) prepared freshly with 940 nL of Lysis Buffer 1 (1 μL of 10X reaction buffer is diluted in 2.75 μL of water) (SMART-Seq v4 Ultra Low input RNA kit; Takara Bio Inc.) per well, aliquoted with the Mosquito HV (STP Labtech) liquid handling robot. The Reaction buffer was prepared following the manufacturer’s instruction adding 1 μL of RNAse Inhibitor in 19 μL of 10X Lysis Buffer.

We ensured maximum sorting accuracy into the wells of the 384-well plate, using a colorimetric assay with tetramethyl benzidine substrate (TMB, BioLegend, Ref. 421501) and 50 μg/mL of Horseradish Peroxidase (HRP, Life Technologies, Ref. 31490) (Rodrigues and Monard 2016). In the plate layout, we sorted nuclei from SC diet-fed animals in half of the 384-well plate and nuclei from HFHS diet-fed animals in the remaining half. After sorting, every plate was firmly sealed (MicroAmp Thermo Seal lid, #AB0558), shortly vortexed for 10 s, centrifuged (4°C, 2000 x *g* for 1 min), flash-frozen on dry ice, and stored at −80°C, until cDNA synthesis. A total of four 384-well plates were sorted for this study.

### snRNA-seq2

The single-nucleusRNA-seq2 methodology was used to capture a high number of transcripts from frozen tissues, allowing for the generation of double-stranded full-length cDNA as described and detailed by Richter *et al*. (Richter et al. 2021) In brief, the reaction volumes were miniaturized with the aid of the Mosquito HV robot. Per well, 2190 nL of Lysis Buffer 2 (LB2) were dispensed. The final volume of the mixture of Lysis Buffer (LB1 and LB2) was 3.125 μL, containing NP40 2%, Triton-X100 1%, 1/300,000 diluted ERCC RNA spike-in, 3′ SMART-seq CDS Primer II A and RNAse-free water.

Every flash frozen sorted plate was thawed directly on a −20 °C chilled metallic holder while LB2 was added by the Mosquito HV robot. The plate was immediately sealed, vortexed 20 s at 2000 rpm, centrifuged at 2000 x *g* for 30 s at 4ºC and placed in a 72 °C for 6 min. ERCC spike-ins (Thermo Fischer Scientific, Ref. 4456740; Lot num 00892098) were diluted 1 in 10, with RNAse-free water with 0.4 U/μL Recombinant RNase Inhibitor (Takara Clontech, Ref. 2313A) and a fresh dilution of 1 in 300,000 was prepared before the first strand synthesis.

Reverse transcription and Pre-PCR amplification steps were followed as described by the manufacturer with four-times reduced volumes for all steps. The PCR program for the cDNA amplification was performed in a total of 21 cycles of: 1 min at 95 °C, [20 s at 95 °C, 4 min at 58 °C, 6 min at 68 °C] × 5, [20 s at 95 °C, 30 s at 64 °C, 6 min at 68 °C] × 9, [30 s at 95 °C, 30 s at 64 °C, 7 min at 68 °C] × 7, 10 min at 72 °C. After cDNA synthesis, the yield was assessed in an Agilent Bioanalyzer with a High Sensitivity DNA kit.

### Library preparation for snRNA-seq2 and sequencing

Sequencing libraries were prepared using the Illumina Nextera XT DNA Sample Preparation kit (Illumina, Ref. FC-131-1096) and the combination of 384 Combinatorial Dual Indexes (Illumina-Set A to D, Ref. FC-131-2001 to FC-131-2004). Using the Mosquito HV robot, the reaction volumes of the Nextera XT chemistry were miniaturized, and the steps followed minutely as described by Richter et al. 2021 (Richter et al. 2021; Mora-Castilla et al. 2016). In brief, 500 nL of the undiluted cDNA were transferred to a new 384 well-plate containing 1500 nL of Tagmentation Mix (TD and ATM reagents). Accordingly, all Nextera XT reagents (NT, NPM and i5/i7 indexes) were added stepwise to a final library volume of 5 μL per well. The final PCR amplification was performed through 12 cycles. Once the libraries were prepared, 500 nL from each well were pooled together into a tube (total volume of ∼192 μL) to perform a final AMPure XP bead (Beckman Coulter, Ref. A63882) clean-up step. For our single nuclei libraries, two consecutive clean-ups with a ratio of sample to bead 0,9X led to library sizes between 200 and 1000 bp. The final libraries were assessed using a HS DNA kit in the Agilent Bioanalyzer, and prior to sequencing, the libraries were quantified using a Collibri library quantification kit (Thermo Fischer Scientific, Ref. A38524100) in a QuantStudio 6 Flex (Life Technologies) for higher accuracy. Each plate, counting with a total of 384 libraries, was pooled together into one final library. A total of 4 final libraries were sequenced using an Illumina NovaSeq 6000 NGS sequencer in an SP XP flowcell, in a paired-end 150 bases length. Sequencing was performed at the Helmholtz Zentrum München (HMGU) by the NGS-Core Facility.

### snRNA-seq2 analysis

The snRNA-Seq2 pipeline used in this study was generated using *nextflow* (Di Tommaso et al. 2017). In the first step, technical replicates of the samples were merged, and reads were then mapped to rRNA (from Ensembl, GRCm38 release 102) using Bowtie2 (version 2.3.4.3, (Langmead and Salzberg 2012)) and subsequently, unmapped reads were mapped to the mm10 genome (from Ensembl, GRCm38 release 102) using STAR (2.7.0d, (Dobin et al. 2013)). Count matrices were generated by counting reads corresponding to gene exons using featureCounts (version 1.6.3, (Liao, Smyth, and Shi 2014)). Counts were transformed to FPKM and TPM using the actual feature length as described by Wagner et al. (Wagner, Kin, and Lynch 2012). The single-nuclei transcriptomics analysis of raw count matrix loaded into python and stored as AnnData object was performed using Scanpy version 1.7.1 (Wolf, Angerer, and Theis 2018). Filtering of the cells was based on a minimum count number of 250. Filtering was applied to the matrix of 1536 single nuclei and 55579 genes. Nuclei that had >1000 and <4500 detected genes were kept. Genes detected in fewer than 25 cells and with read count below 250 were filtered out. Hence, only the nuclei having less than 4500 genes detected and a library size between 10000 and 200000 reads were kept. Applying this filtering strategy, the final matrix comprises 1202 single nuclei and 13867 genes. The remaining cell vectors were normalized using the R-package *scran* (Lun, McCarthy, and Marioni 2016) in the default setting, employing the ERCC spik-ins. Batch effect corrections were performed employing *combat* (Johnson, Li, and Rabinovic 2007) using plates as a covariate. To generate UMAP plots, we took the top 15 PCs and used the PC space to compute a k-nearest neighbor (kNN) graph (k=50, metho=umap). *Leiden* clustering (resolution=0.5, flavor=vtraag) was computed based on the kNN graph. Differential gene expression analysis compared the groups of interest using Welch’s t-test while applying Benjamini-Hochberg for multiplicity correction. In order to identify transcript intersections as well as their aggregates at single-cell resolution, the novel *UpSet* visualization technique was applied according to the previously published protocol (Lex et al. 2014) using R-package *UpSet* (Conway, Lex, and Gehlenborg 2017).

### Statistics

Data analysis was conducted using GraphPad Prism (Version 5). Normally distributed data were analyzed by student’s T-test or one- or two-way analysis of variance (ANOVA) with Bonferroni or Tukey post-hoc analyses to determine statically significant differences. Data were screened using the maximum normal residual Grubb’s test to screen for singular, statistically significant outliers. P-values ≤ 0.05 were considered statistically significant. All data are presented as mean ± standard error (SEM).

## Figures and figure legends

**Figure S1. Related to Figure 1:**
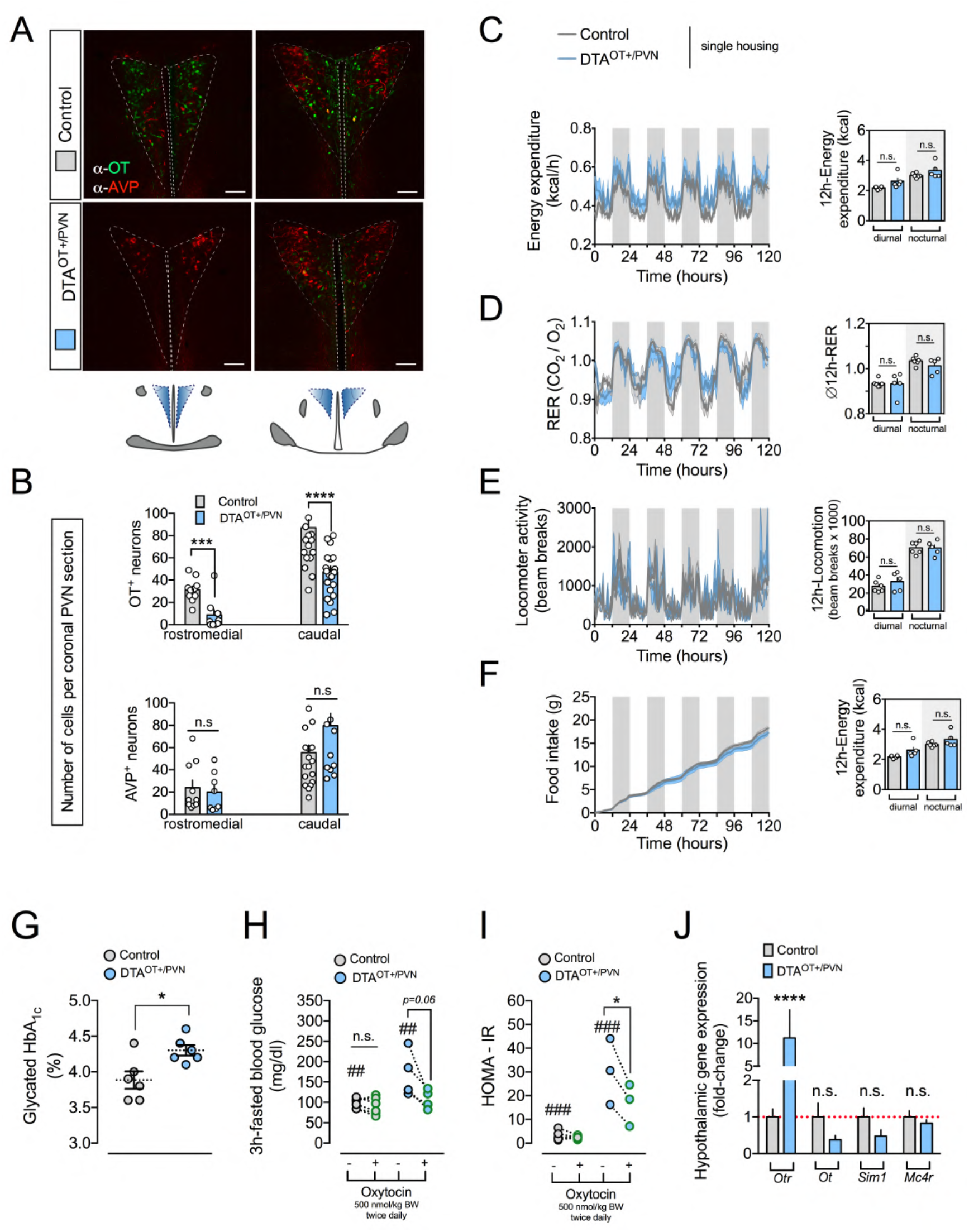
Virus-mediated ablation of PVN^OT^ neurons induces hyperphagic obesity that is rectifiable by exogenous oxytocin treatment and associated with CCK resistance. **(A)** Representative confocal micrographs of brain sections showing OT^+^ neurons (green) and AVP^+^ neurons (red) of control mice and DTA^OT+/PVN^ mice at the rostromedial and caudal levels of the PVN. Scale bar, 100 μm. **(B)** Corresponding quantification of OT neuron count (upper panel) and AVP neuron count (lower panel) control mice and DTA^OT+/PVN^ mice at the rostromedial and caudal levels. n = 5-7, 3-5 hemisections per mouse. **(C)** Hourly energy expenditure as measured by indirect calorimetry in metabolic cages of single-housed control mice and DTA^OT+/PVN^ mice (left panel) as well as average 12h-energy expenditure (right panel). Data are presented as mean ± SEM. ** P < 0.01, **** P < 0.0001. n = 5-7 mice (two-way ANOVA (left panel) and unpaired Student’s *t*-test (right panel). **(D)** Hourly respiratory exchange ratio (RER) as measured by indirect calorimetry in metabolic cages of single-housed control mice and DTA^OT+/PVN^ mice (left panel) as well as average 12h-RER (right panel). Data are presented as mean ± SEM. n.s., not significant. n = 5-7 mice (two-way ANOVA (left panel) and unpaired Student’s *t*-test (right panel)). **(E)** Hourly locomotor activity as measured by beam breaks in metabolic cages of single-housed control mice and DTA^OT+/PVN^ mice (left panel) as well as average 12h-locomotion (right panel). Data are presented as mean ± SEM. n.s., not significant. n = 5-7 mice (two-way ANOVA (left panel) and unpaired Student’s *t*-test (right panel)). **(F)** Cumulative food intake of single-housed control mice and DTA^OT+/PVN^ mice (left panel) as well as average 12h-food intake (right panel). Data are presented as mean ± SEM. n.s., not significant. n = 5-7 mice (two-way ANOVA (left panel) and unpaired Student’s *t*-test (right panel)). **(G)** Quantification of glycated HbA_1C_in a separate cohort of DTA^OT+/PVN^ mice and control mice. Data are presented as mean ± SEM. n.s., not significant. * P < 0.05. n = 6 mice (unpaired Student’s *t*-test). **(H)** Quantification of 3h-fasted blood glucose before and after treatment with bi-daily OT (500 nmol/kg BW; *s*.*c*.) in DTA^OT+/PVN^ mice and control mice. Data are presented as mean ± SEM. ## P < 0.01, n.s., not significant. n = 5-7 mice (one-way ANOVA and paired Student’s *t*-test). **(I**) Quantification of HOMA-IR before and after treatment with bi-daily OT (500 nmol/kg BW; *s*.*c*.) in DTA^OT+/PVN^ mice and control mice. Data are presented as mean ± SEM. * P < 0.05, ### P < 0.001, n.s., not significant. n = 5-7 mice (one-way ANOVA and paired Student’s *t*-test). **(J)** Relative gene expression of mRNA for *Otr, Ot, Sim1* and *Mc4r* in the hypothalamus of DTA^OT+/PVN^ mice normalized to control mice. Data are presented as mean ± SEM. * P < 0.05, n.s., not significant. N = 4 mice (unpaired Student’s *t*-test).

**Figure S2. Related to Figure 2:**
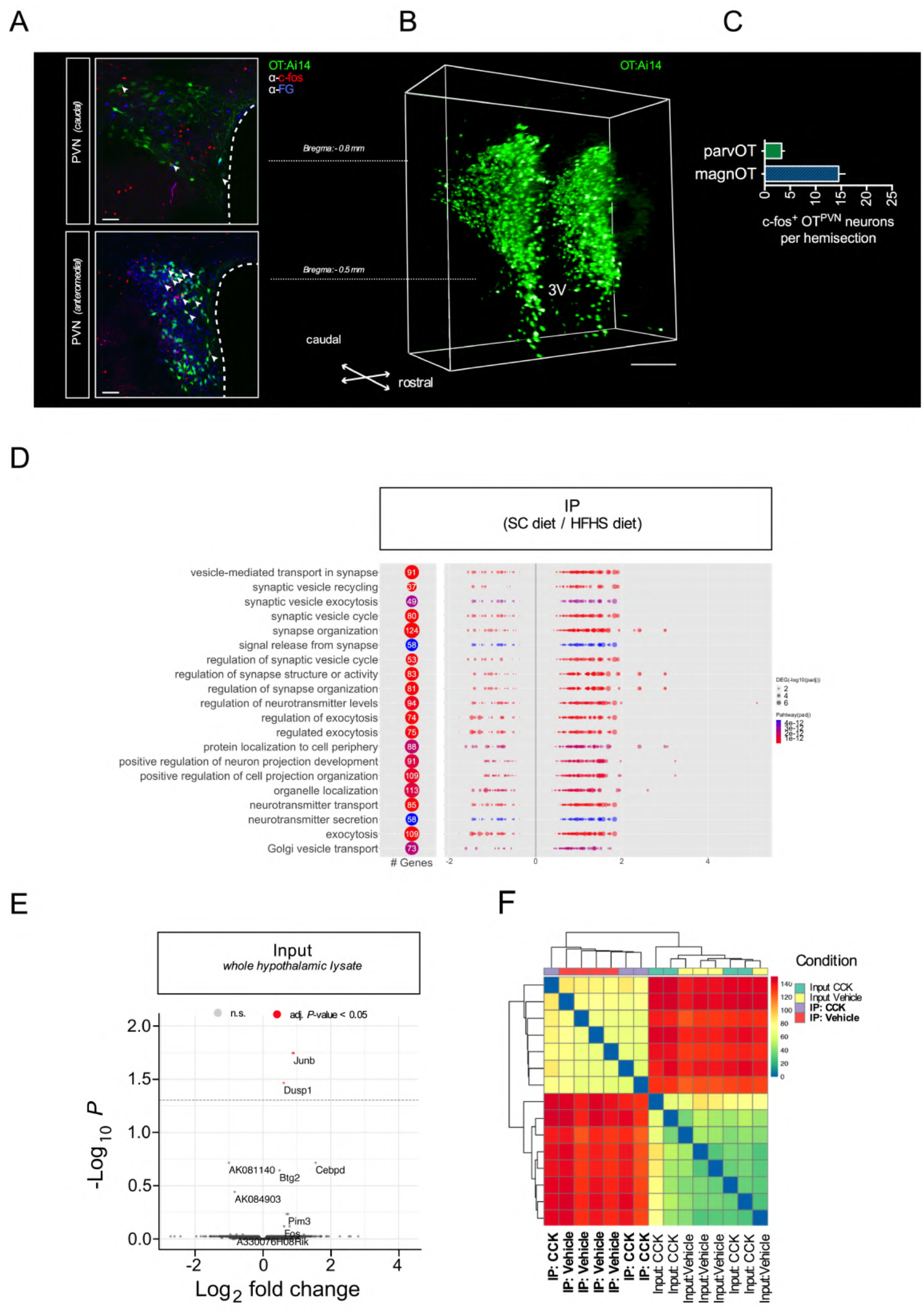
Chronic exposure to a HFHS diet impairs the electrical and transcriptional activation f PVN^OT^ neurons in response to peripheral CCK. **(A)** Representative confocal micrographs of coronal brain sections from adult male OT:Ai14 reporter mice containing the PVN at the caudal (upper panel) and anteromedial (lower panel) level relative to bregma. Mice received fluorogold (FG; 15 mg/kg BW *i*.*p*.) 7 days prior sacrifice in order to label magnOT neurons, which form neurohemal contacts at the posterior pituitary (FG^+^; blue). On the day of experiment, mice were injected with CCK (20 μg/kg BW *i*.*p*.) and consequent activation of PVN^OT^ neurons (green) was quantified by means of nuclear c-fos immunoreactivity (red). Scale bar, 50 μm. **(B)** 3D rendered confocal scan of iDISCO-cleared coronal brain section from an adult male *OT:Ai14* reporter mouse spanning the entirety of the PVN (1 mm). Scale bar, 1 mm. **(C)** Quantification of total c-fos^+^ PVN^OT^ neuronal subpopulations from (A) differentiating between parvOT (FG^-^) and magnOT (FG^+^) subsets. **(D)** GO enrichment analysis of DEG comparing IP of OT:RiboTag mice either fed SC diet or HFHS diet. Top enriched pathways number of DEG are indicated in the left panel, while the color indicates the adjusted p-value. Each pathway DEG are represented as dots, and plotted against log-fold changes, while the size indicates the adjusted p-values. **(E)** Volcano plot highlighting the DEG in the input from OT:RiboTag mice fed SC diet receiving CCK (20 μg/kg BW *i*.*p*.) relative to vehicle. **(F)** Heat map of sample-to-sample distance matrix for overall normalized gene expression read counts of both input and IP samples of OT:RiboTag mice fed either SC diet or HFHS diet that were additionally treated with either CCK (20 μg/kg BW *i*.*p*.) or vehicle. Euclidean distance clustering dendrograms are displayed above.

**Figure S3. Related to Figure 3:**
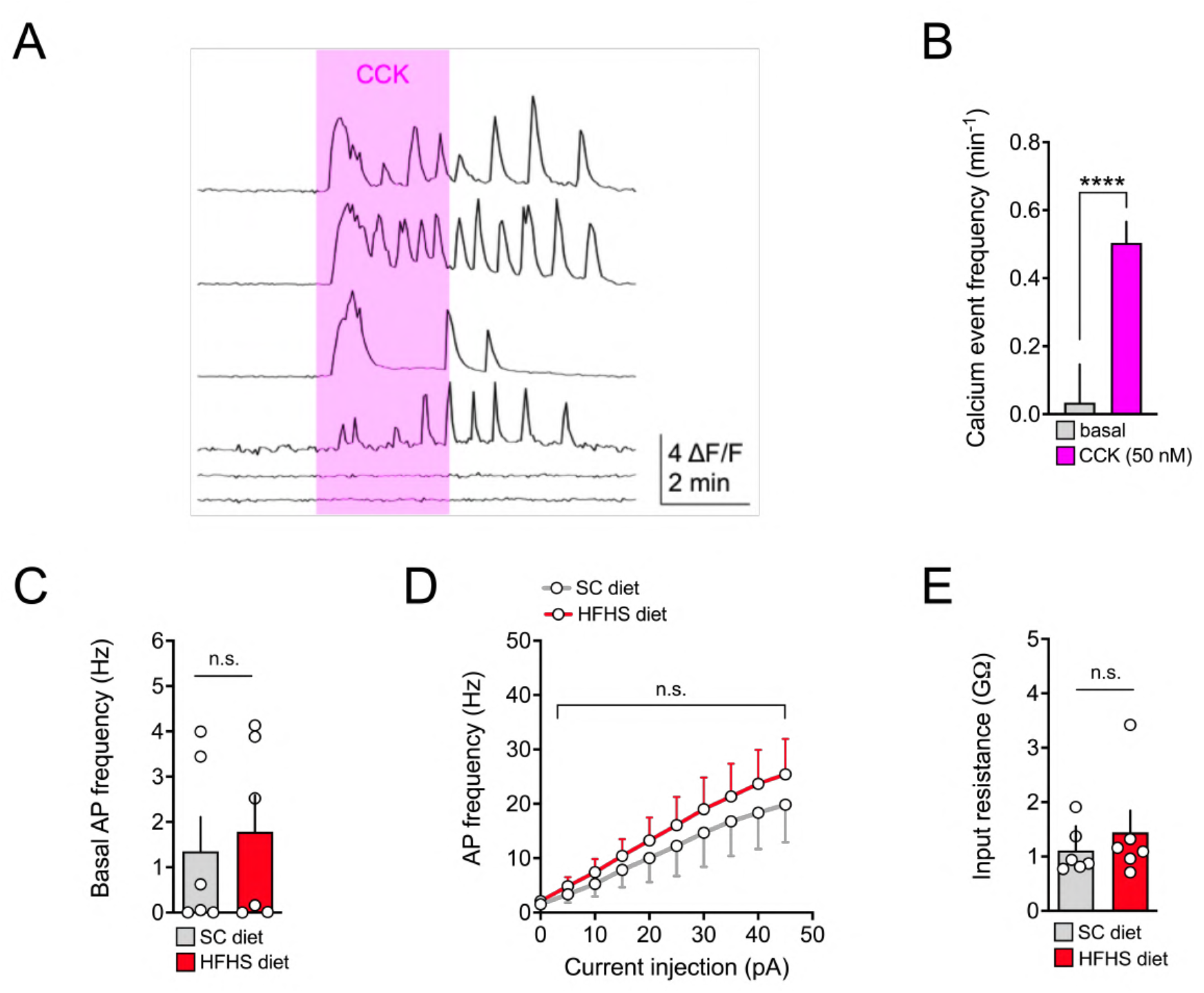
PVN^OT^ neurons are activated by CCK *via* a direct, CCK_A_R-dependent mechanism in lean but not obese mice. **(A)** Cytosolic Ca^2+^ transients of individual PVN^OT^ neurons (lower panel) upon bath application of CCK (50 nM) in the presence of synaptic blockers. **(B)** Quantification of Ca^2+^ event frequency as summary data of all imaged neurons. Data are presented as mean ± SEM. **** P < 0.0001. n = 1 mouse, 49 neurons (unpaired Student’s *t*-test). **(C)** Quantification of basal action potential frequency of putative magnOT neurons. Data are presented as mean ± SEM. n.s. = not significant. n = 2-3 mice/ 6 neurons per mouse (unpaired Student’s *t*-test). **(D)** Quantification of firing frequency as a function of injected current of putative magnOT neurons. Data are presented as mean ± SEM. n.s. = not significant. n = 2-3 mice/ 6 neurons per mouse (unpaired Student’s *t*-test). **(E)** Quantification of input resistance of putative magnOT neurons. Data is represented as mean ± SEM. n.s. = not significant. n = 2-3 mice/ 6 neurons per mouse (unpaired Student’s *t*-test).

**Figure S4. Related to Figure 4:**
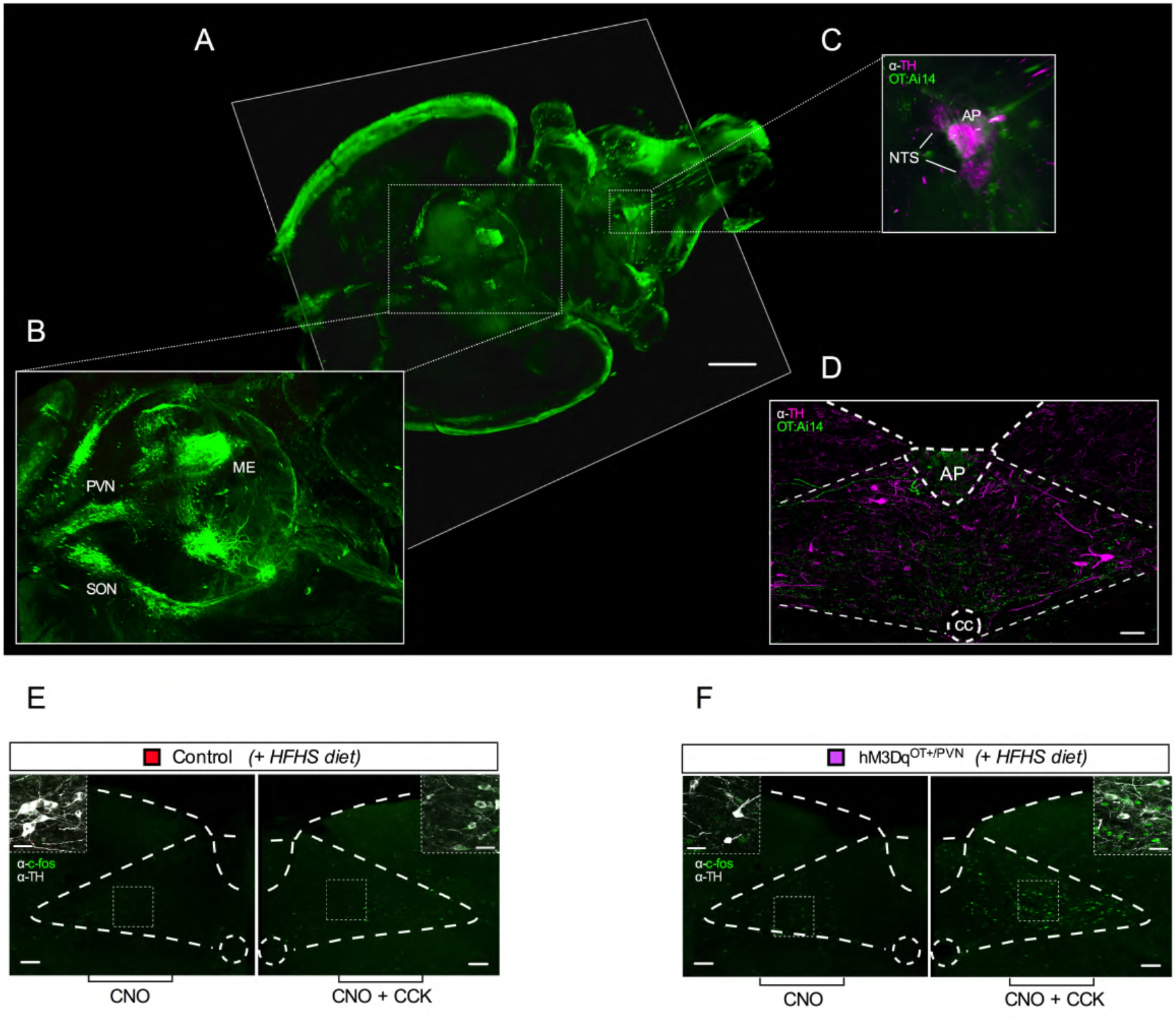
Blunted suppression of food intake in response to CCK on a HFHS diet is reinstated by concomitant chemogenetic activation of PVN^OT^ neurons. **(A)** 3D whole-brain image (horizontal view) of an iDISCO-cleared OT:Ai14 reporter mouse brain subjected to light-sheet fluorescence microscopy. Scale bar, 1 mm. **(B)** 3D rendered zoom-in image (dashed line insert) of the hypothalamus showing the anatomical organization of the OT system. **(C)** 3D rendered zoom-in image (dashed line insert) of the dorsal vagal complex (NTS and AP) in the brainstem containing catecholaminergic TH^+^ neurons (magenta) and its innervation by OTergic fibres (green). **(D)** Confocal micrograph of a coronal brain section of the NTS displaying catecholaminergic TH^+^ neurons (magenta) and their innervation by OTergic fibres (green) at high resolution. Scale bar, Scale bar, 100 μm. **(E)** Representative confocal micrographs of NTS brain sections from HFHS diet-fed control mice showing c-fos immunoreactive cells (green) following either CNO or CNO+CCK; inserts displaying the extent of co-localization with TH (gray). Scale bar, 100 μm and 20 μm (insert). **(F)** Representative confocal micrographs of NTS brain sections from HFHS diet-fed hM3Dq^OT+/PVN^ mice showing c-fos immunoreactive cells (green) following either CNO or CNO+CCK; inserts displaying the extent of co-localization with TH (gray). Scale bar, 50 μm and 20 μm (insert).

**Figure S5. Related to Figure 5.**
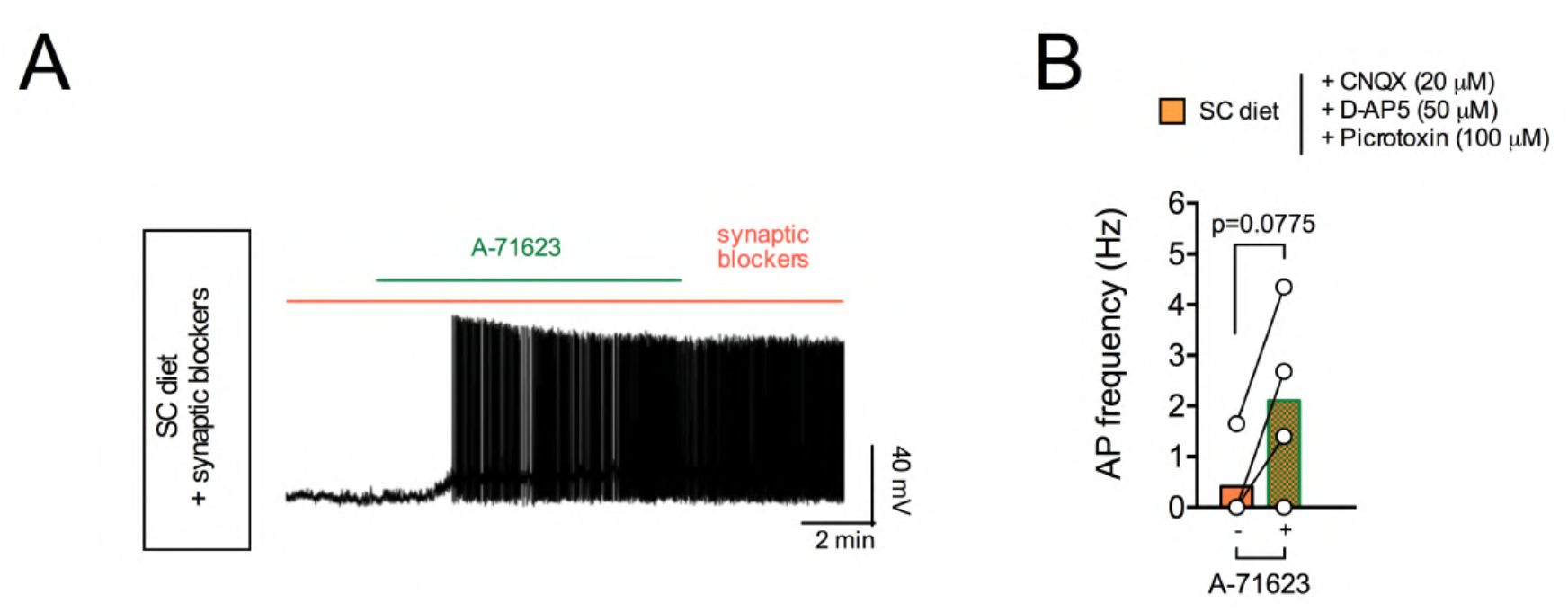
Intersectional regulation of hypothalamic OT neurons by CCK_A_R and κ-opioid receptors is dependent on dietary context. **(A)** Representative traces of action potential frequency of magnOT neurons derived from adult male *OT:Ai14* reporter mice fed SC diet in response to bath-applied A-71623 (25 nM) pre-treated with synaptic blocker. **(B)** Summary of changes in action potential frequency (right panel). Data are presented before and after application of A-71623 as mean ± SEM. n = 1 mouse/ 3 neurons per mouse (paired Students *t*-test).

